# The bacterial cell membrane acts as a dynamic heme reservoir during group B *Streptococcus* bloodstream infection

**DOI:** 10.64898/2026.05.15.725516

**Authors:** GH Hillebrand, HA Stephenson, EJ Giacobe, AS Neel, SM Carlin, FD Kemp, TA Hooven

## Abstract

During bloodstream infection, most bacterial pathogens maintain homeostatic levels of heme, which serves as an essential biochemical cofactor and iron source, but becomes toxic at high intracellular concentrations. Well-characterized, surface exposed heme binding and acquisition systems exist in several blood-borne bacterial species. However, some gram-positive bacteria that invade the bloodstream do not encode surface displayed heme acquisition systems, despite showing clear evidence of heme utilization in blood. An example is *Streptococcus agalactiae* (group B *Streptococcus;* GBS), which is a major cause of infection in neonatal and immunocompromised populations. Here we show that GBS uses its cell membrane as a dynamic heme reservoir, which functions as the primary site of environmental heme capture, sensing, and transmembrane flux. Using positive and negative genetic selection screens, targeted mutagenesis, membrane fractionation, and spectroscopic heme detection and binding assays, we demonstrate that heme is partitioned into the GBS cell membrane, where it is sensed by the histidine kinase HssS and extracted for intracellular use by the CydDC transporter. Genetically disrupting the function of either HssS heme sensing or CydDC membrane heme extraction attenuates bacterial survival in human whole blood and in a mouse model of bacteremia. These results suggest that cell membrane-localized heme homeostasis is a determinant of fitness during blood survival. This work expands the current models of bacterial heme physiology and provides evidence that membrane localized, homeostatic heme reservoirs may represent an underrecognized strategy for blood-borne pathogens that lack canonical heme acquisition systems.

## Background

Bacteria invading the bloodstream face significant chemical and metabolic challenges, including nutrient acquisition, energy production, and resisting oxidative damage ^1^. Heme, which consists of protoporphyrin IX in complex with ferrous iron—an essential biochemical cofactor—is a key participant in redox reactions across the tree of life, including controlled electron transport that can generate ATP in respiration-enabled bacteria ^1–3^. Paradoxically, accumulation of excess heme subjects bacteria to destructive oxidative stress, necessitating sensing and export systems to maintain a sustainable intracellular milieu ^4,5^.

Accordingly, some bacteria have surface enzyme networks that sense extracellular heme, extract it from sequestering host hemoproteins, and transport it from the extracellular to intracellular spaces, where it serves as an iron source, substrate for respiration, and an exportable electron sink to detoxify reactive oxygen species ^6–12^. Other bacteria encode endogenous heme synthesis pathways, enabling a ready supply ^2,13^. Despite substantial progress in defining systems for bacterial heme acquisition, utilization, and detoxification, most existing models implicitly treat heme as either extracellular—requiring surface capture and transport—or cytoplasmic, where it is incorporated into metabolic processes or exported to prevent toxicity.

These models, however, do not account for bacteria that utilize heme during bloodstream invasion but neither synthesize it nor express surface-displayed heme binding proteins to extract the molecule from the surrounding environment. An example is group B *Streptococcus* (*Streptococcus agalactiae*, GBS), an encapsulated gram-positive pathobiont with virulence properties that manifest primarily during the perinatal period or in immunocompromised hosts ^14–17^.

Genetic and prior experimental evidence indicates that heme promotes GBS bacteremia. The terminal oxidoreductase composed of protein subunits CydA and CydB, which together form the cytochrome bd oxidase, are highly conserved among GBS strains, encoded by the first two genes in the *cydABCD* operon, and have been shown to enable heme-based respiratory metabolism ^18^. Two heme export pathways in GBS have also been described. The first, HrtAB, is a well-characterized heme transporter with orthologs in *Staphylococcus aureus* ^5,19^, *Lactococcus lactis* ^20,21^, and *Bacillus anthracis* ^22^. A second export system, encoded by distinct *pefAB* and *pefCD* gene pairs, is expressed in GBS and regulated by an intracellular protoporphyrin sensor PefR ^23^.

As in several orthologous systems, HrtAB activity in GBS is regulated by the heme sensing system (HssRS) two-component system ^22,24^. Unlike other bacterial pathogens, GBS encodes *hssRS* on a polycistronic operon with *hrtAB* ^25^. One of the best-characterized HssS structures, from *S. aureus*, has a conformationally complex extracellular domain, which has been shown to participate in forming a heme binding pocket at its junction with the cell membrane bilayer’s outer lipid leaflet. The structural basis of GBS HssS signal transduction has not been examined, nor is it understood how heme is shuttled from the extracellular milieu by GBS to utilize it in cellular processes.

Therefore, despite the clear contribution of heme to GBS survival during bloodstream invasion, open questions remain about how heme is obtained and processed. Here we examine mechanisms of heme acquisition and utilization, using negative and positive library selection screens, confirmatory targeted mutations, and *in vitro*, *ex vivo*, and *in vivo* models of bacteremia. We show that porphyrin partitioning into the cell membrane lipid bilayer establishes a reservoir of heme that can be directly sensed through a novel inner membrane leaflet heme-binding pocket on HssS and transported from the intramembrane space through a laterally loaded CydDC importer previously described in *Escherichia coli*.

Disruption of membrane heme homeostasis in GBS attenuates survival in blood and impairs virulence during systemic infection. Our findings advance the concept of membranous deposition as a mechanism by which bacteria invading the bloodstream can store heme for import and metabolic use without necessitating active harvesting from the environment, and connect two bacterial heme-protein interactions that thermostatically regulate the intramembrane heme pool. This work broadens our understanding of how invading bacteria persist and adapt during bloodstream infection and points to new potential targets for disrupting bacterial infections that rely on host heme for persistence and spread.

## Results

### HssRS, which lacks an externalized heme binding site, is the sole heme-responsive GBS two-component system

22 distinct two-component systems (TCS) have been identified in GBS. While not all are encoded by every GBS strain, most are highly conserved ^26^. TCS are crucial bacterial sense-and-response signal transduction systems that modulate transcription to promote adaptive fitness to environmental fluctuation. The fact that GBS encodes 22 TCS—considerably more than closely related pathobionts such as *Streptococcus pneumoniae* and *Streptococcus pyogenes*, which each encode 13—has been interpreted as evidence of GBS evolution to survive across diverse environmental conditions ^26^.

One GBS TCS, the HssRS system, has previously been identified as responsive to heme exposure ^25^. The GBS *hssRS* genes are expressed on an operon that also contains the *hrtAB* genes encoding the heterodimeric HrtAB heme exporter. This fact, along with partial homology to the *S. aureus* HssRS TCS, prompted prior studies that convincingly demonstrated HssRS conditional essentiality during exposure to elevated heme concentrations. The same work also showed that heme signaling through the HssRS system led to upregulation of the entire *hrtAB*-*hssRS* operon ^25^.

Most of the signals that modulate the 22 TCS encoded by GBS are unknown, and we considered it plausible that supplementary systems beyond HssRS might be involved in heme homeostasis. To test this possibility, we performed a negative selection screen for susceptibility to heme toxicity on a library of CRISPR interference (CRISPRi) TCS variants in the GBS serotype Ia background strain A909 modified to express catalytically inactive Cas9 (A909 *dcas9*) ^27^. The library was composed of TCS knockdown strains (see **Suppl. Fig. 1** for gene targets) that were pooled together in equal colony forming unit (CFU) concentrations. Plasmid DNA was purified from the CRISPRi library replicates before outgrowth (input samples), then the library was grown in nonselective tryptic soy broth (TSB) with or without overnight exposure to supplemental 10 μM hemin chloride (hereafter referred to as hemin). After the control or experimental exposure, plasmid DNA from the mixed library was again collected (output samples). Targeted amplicon sequencing of the CRISPRi protospacer insertion site was performed and protospacer read counts were used to determine fractional contribution of each GBS variant to the total population. Normalized protospacer sequence counts were then compared across input and output samples, allowing derivation of fold change values. To isolate hemin specific effects on fitness, fold change under hemin exposure was adjusted by fold change under the control TSB condition to derive Δfold change.

Each of the 22 TCS expressed by A909 was separately targeted with two sgRNA sequences designed to disrupt the first gene in the polycistronic sensor/response operon (**Suppl. Data 1**). Unnamed TCS were numbered by their order of appearance in the A909 genome; previously named TCS were labeled accordingly ^26^. Two intended knockdown strains targeting the essential TCS VicRK ^28^ were nonviable and were not included in the screen. Additionally, one of the two knockdown strains for the FspRS and LtdRS systems did not generate protospacer reads from the input sample, possibly due to PCR failure across those two protospacers; we therefore only used the successfully sequenced protospacer as a marker for knockdowns of these two TCS. This resulted in a total of 40 knockdown strains included in the library.

We performed the library screen on three independent replicates, which were sequenced with different multiplexing barcodes. Reproducibility across the replicates was high, as shown in the principal component analysis of z-score adjusted input and output read counts plotted in **Fig. 1A**, with each library screen replicate showing similar fitness responses across TSB and hemin-exposure conditions. Dual targeting protospacers had congruent effects, with the majority clustering around a Δfold change value of zero, indicating no effect on fitness in the presence of hemin exposure (**Fig. 1B** and **Suppl. Fig. 1**). However, the library screen revealed a pronounced heme-induced fitness defect, congruent across both targeting protospacers, in the HssRS knockdowns, with a mean 18.33-fold reduction in relative abundance compared to the TSB control (**Fig. 1B-C**). By contrast, the next most inhibited library constituent (targeting the RgfAC system) had a 1.96-fold reduction in hemin compared to TSB (**Fig. 1C**).

**Figure 1:**
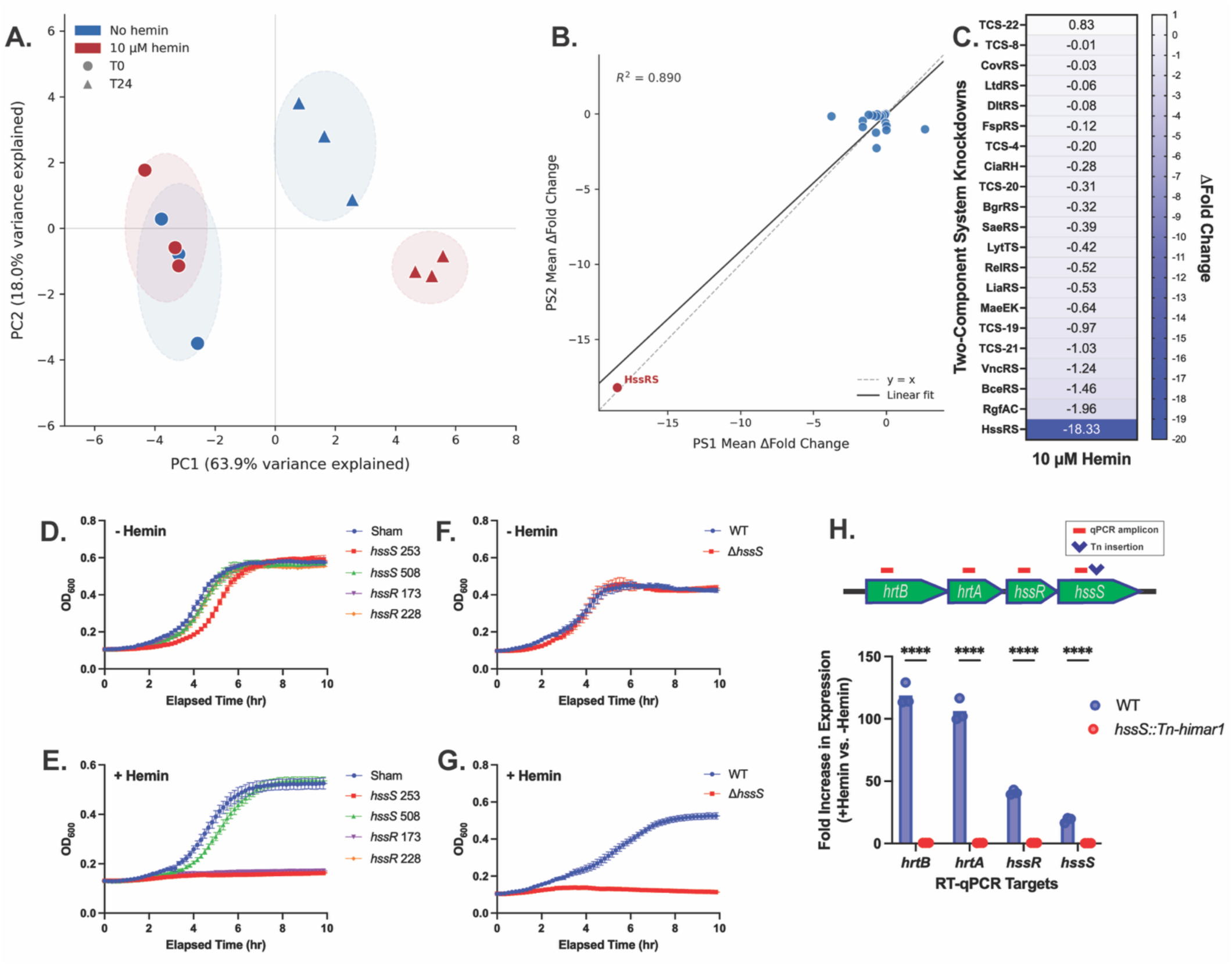
HssRS is the sole GBS TCS responsive to heme exposure, necessary for resisting heme toxicity through autoregulated transcriptional upregulation of the HrtAB exporter. A CRISPRi library with knockdowns of all two-component signal transduction systems in GBS strain A909 *dCas9* was screened in TSB and TSB supplemented with 10 μM hemin. Fitness was determined from relative quantification of targeting protospacers in the input and output libraries. Three independent biological replicates of the same CRISPRi library were tested. Principal component analysis of the three biological replicates’ selection characteristics in TSB and hemin showed clustering by growth condition (**A**). A correlation plot shows mean Δfold change for two corresponding protospacers (PS) for the dual-targeted TCS across three biological replicates, along with a linear best fit (R^2^=0.89) and x=y reference line (**B**). Fitness of each tested TCS displayed as mean Δfold change among targeting protospacers within the library population, comparing hemin exposure to TSB (**C**). Individual A909 CRISPRi knockdown strains and a sham-targeted negative control strain were used to generate growth curves in TS media or TS media supplemented with 10 μM hemin (**D-E**, n=3, error bars show SEM). WT and a Δ*hssS* targeted deletion strain were used to generate growth curves in 5 μM heme (**F-G**, n=3, error bars show standard error of the mean). WT A909 and a transposon mutant strain in which *hssS* is interrupted were used to measure expression across the *hrtBA-hssRS* operon in hemin-negative and hemin-positive growth conditions (**H**, **** p<0.001, ANOVA with Bonferroni correction).

To validate the CRISPRi results, we individually tested four HssRS CRISPRi strains—the two *hssR* knockdowns that were used in the CRISPRi screen and two additional *hssS* knockdowns. As expected, when exposed to 10 μM hemin, these strains showed pronounced growth defects. The growth phenotype was most significant in the *hssR* 173, *hssR* 228, and *hssS* 253 strains; it was less severe but still present in the *hssS* 508 strain, where the CRISPRi sgRNA target is near the 3’ end of the gene, potentially attenuating its effects (**Figs. 1D-E**). Next, we used Cas12a-mediated mutagenesis to create a targeted deletion strain lacking *hssS* (Δ*hssS*), which showed growth defects under hemin stress conditions **(Fig. 1F-G**).

RT-qPCR on WT and a transposon insertion mutant in which *hssS* is interrupted by a Himar1 mini-transposon downstream of the *hssS* qPCR probe pair (A909 *hssS*::*Tn-Himar1*) ^29^ demonstrated that expression of the TCS and heme export system was significantly increased during hemin exposure in a *hssS*-dependent manner, indicating autoregulation of the entire *hrtAB*-*hssRS* operon by the HssRS system (**Fig. 1H**). To examine whether the HssRS-HrtAB signaling axis was due to heme reception specifically or was a nonspecific effect of membrane stress, we subjected WT GBS to nonlethal concentrations of chemical and pharmacologic membrane disruptors (triton, ethanol, H_2_O_2_, ampicillin), testing HrtAB responses through RT-qPCR. Detergent treatment with 0.01% Triton caused modest, variable HrtAB induction, but none of the tested membrane stressors had a HrtAB signaling response of the magnitude caused by hemin, indicating that hemin is the specific molecular signal sensed by HssS (**Suppl. Fig. 2**).

Together, these results showed that HssRS is the sole GBS TCS responsible for environmental heme sensing, that autoregulation of its own operon (together with genes encoding HrtAB) is a reliable downstream readout of its HssRS activity, and that HssRS function is essential for GBS survival in heme-rich environments. These were crucial premises to establish for the further mechanistic studies of GBS heme utilization and homeostasis described below.

### Targeted domain mutagenesis indicates roles of the HssS intracellular and transmembrane domain 1 in GBS heme sensing

Comparing HssS in GBS to other gram-positive pathobionts demonstrated significant sequence differences, with predicted effects on tertiary and quaternary structures, raising questions about where heme might bind to influence HssS signaling in GBS. Unlike the well-characterized HssS proteins of *B. anthracis* and *S. aureus*, which fold into membrane-inserted homodimers with a complex extracellular structure believed to be involved in heme binding and thus signal induction ^22,24^, the GBS HssS homodimer AlphaFold3 model does not include a substantial extracellular domain. Instead, GBS HssS has a short, 10-residue linker predicted to connect its two transmembrane alpha helices (**Fig. 2A** and **Suppl. Fig. 3**). This rudimentary extracellular structure is not predicted by domain function models to form a recognizable heme binding site.

**Figure 2:**
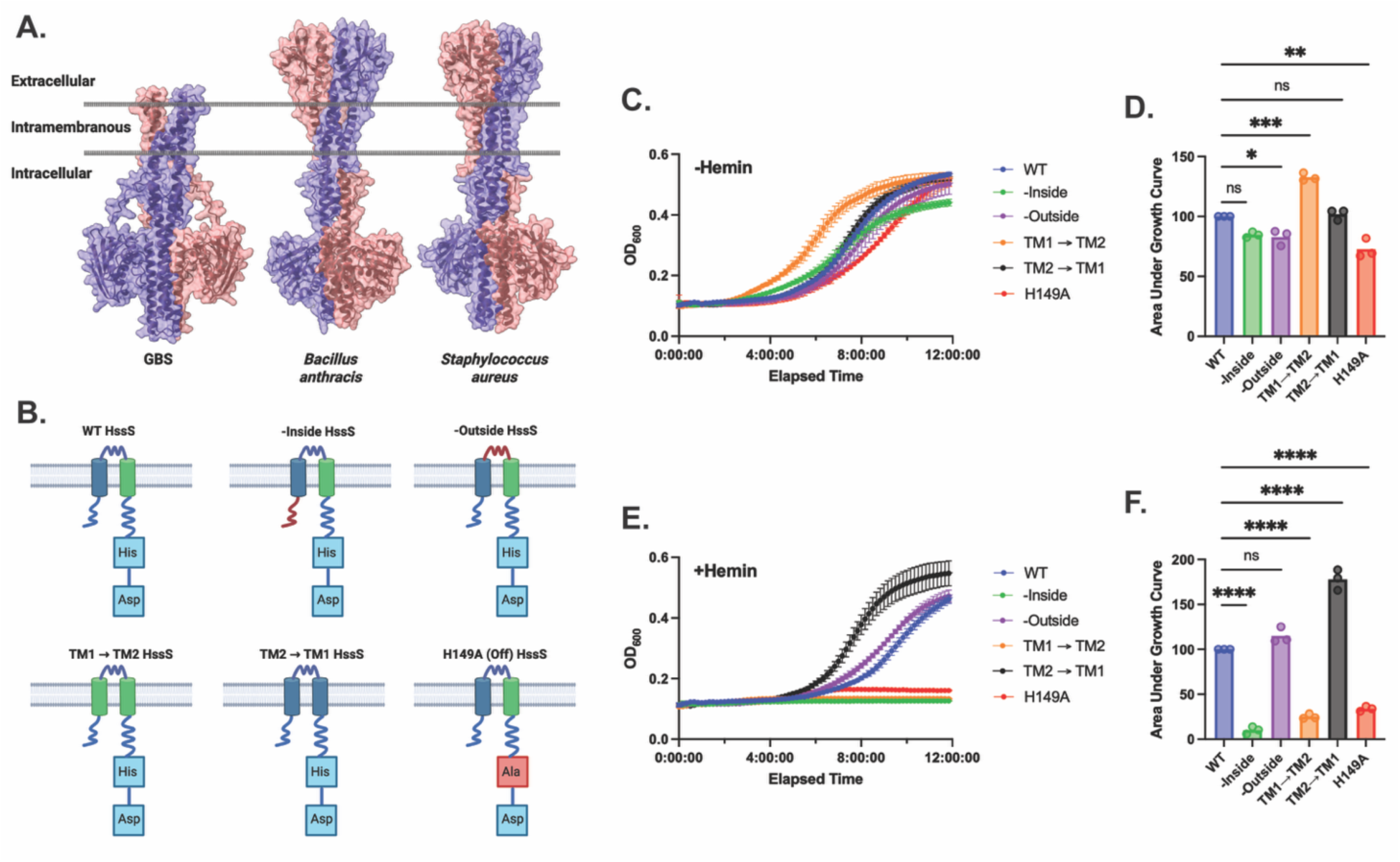
Comparative HssS domain structure analysis and GBS domain swap experiments suggest structure-function diHerences from close orthologs. AlphaFold3 models of GBS, *B. anthracis*, and *S. aureus* HssS protein homodimers with cell membrane lipid bilayers overlaid show that GBS HssS lacks the characteristic extracellular domain of previously characterized examples (**A**). Schematic of domain mutant HssS designs; all domain mutant variants were expressed from plasmid (**B**). Growth curves of WT and domain mutant strains in TS media without hemin supplementation (**C**) with comparisons between areas under the growth curves (**D**). Growth curves of WT and domain mutant strains in TS media with 10 μM hemin supplementation (**E**) with comparisons between areas under the growth curves (**F**). For all growth curves and comparisons, n=3, error bars show standard error of the mean, * p < 0.05, ** p <0.01, *** p < 0.005, **** p < 0.001, ns = nonsignificant, ANOVA with Bonferroni correction.

Given the structural differences between GBS HssS and previously described orthologs, we sought to examine where heme sensing occurs on the GBS variant. As a first approach, we tested domain mutant strains in which we altered either the predicted intracellular N-terminus of the HssS protein, the short extracellular bridge, or either of the two predicted transmembrane domains. To generate these domain mutants, we used either 1) an alanine substitution strategy for the intra- and extracellular domains or 2) a domain swap strategy in which each of the two predicted transmembrane domains were replaced with the other. Of note, the intracellular C-terminus domain connected to transmembrane domain 2 is predicted to form a canonical phosphorelay moiety that is essential for histidine kinase function, and is highly conserved across TCS. We designed an alanine substitution sequence in this domain to abolish phosphorelay function, thereby establishing a signal-dead control strain (H149A) in which signaling activity was expected to be constitutively suppressed. All the tested domain mutants were expressed from an extrachromosomal expression plasmid in a Δ*hssS* background (**Fig. 2B**).

These domain mutant strains showed significant differences compared to WT in resistance to heme stress. When grown in rich media without supplemental hemin, all of the strains showed normal bacterial growth phases, with minor rate differences between them (**Fig. 2C-D**). However, when supplemental hemin was added to rich media, these differences were accentuated, with the inside domain and transmembrane domain 1-deficient (TM1 → TM2) swap mutants showing complete growth failure, comparable to the H149A signal-dead control strain (**Fig. 2E-F).** Notably, the WT to the outside domain mutant comparison was not significant in the hemin supplemented condition, further emphasizing differences between GBS HssS and the other species’ variants, where the extracellular portion of the homodimer is believed to be an important participant in heme binding.

This result was suggestive of a GBS HssS heme binding site localized to an intramembrane or intracellular location but was not definitive, given that changes to one region of a membrane-spanning TCS can have conformational effects on the entire protein that affect function. For this reason, we believed it worthwhile to further probe the heme sensing functionality of the GBS HssS homodimer.

### A positive selection screen of a saturated transposon library exposed to lethal hemin concentrations reveals *cydDC* as a GBS heme importer

Our heme exposure studies so far had involved modulation of extracellular heme, but formal testing of an intramembrane or internalized HssS binding sites would only be possible through experimental manipulation of heme concentrations in at least one of those two bacterial cell compartments. For this reason, we undertook to identify a GBS heme import pathway.

HssRS is understood to upregulate the HrtAB heme export system, a concept confirmed by our qPCR experiments described above, yet it is unknown how heme enters GBS. The close GBS relative, *Streptococcus pyogenes* (group A Streptococcus; GAS), encodes a well-characterized set of heme binding and transport proteins—Shr, Shp, and SiaA-H—on a single operon, regulated by MtsR, that together coordinate surface acquisition and import of heme ^7–9,30,31^. However, no dedicated heme import pathway has been described in GBS.

To identify putative heme importers, we performed a positive selection screen on a previously described saturated transposon mutant library ^28,29^ in the presence of a GBS-lethal hemin concentration (100 μM). The premise of the experiment was that deletion of the heme importer would effectively shield that mutant from the toxic oxidative effects of high-concentration intracellular heme, facilitating identification of the permissive mutant and therefore the import protein(s). Using targeted transposon insertion site PCR and Sanger sequencing, we identified 18 separate transposon insertion sites among the surviving colonies retrievable following growth in high-hemin conditions (**Suppl. Data 2**). We interrogated these transposon insertion sites for plausibility, asking whether the insertions interrupted genes with surface anchored domains, transmembrane regions, or sequence annotations pointing to transporter functionality. Using these criteria to select eight candidate heme acquisition genes (**Fig. 3A**), we next tested the individual transposon mutant strains for decreased intracellular heme concentrations following growth in high-hemin conditions.

**Figure 3:**
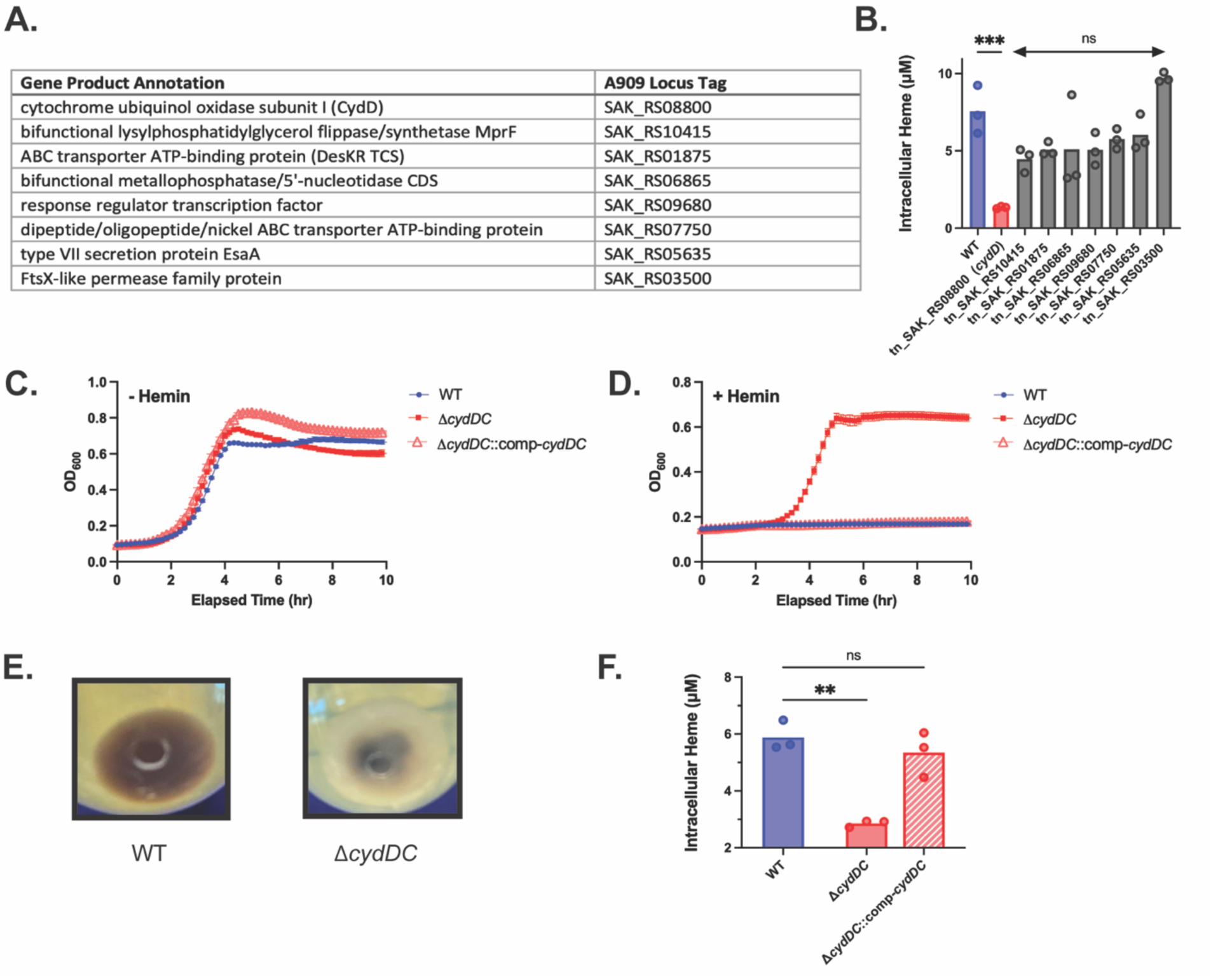
GBS cydDC is a transporter that moves heme from the intramembrane to the intracellular compartment. Candidate heme transporters were selected from a saturated transposon mutant library screen against lethal hemin (100 μM) exposure (**A**). Each candidate transposon mutant was assessed for intracellular heme concentration following growth in supplemented TSB (**B**, *** p < 0.005, ns = nonsignificant, ANOVA with Bonferroni correction). Growth curves of WT, Δ*cydDC*, and Δ*cydDC*::comp-*cydDC* in lethal hemin-negative and lethal hemin-positive conditions (**C-D**, n=3, error bars show standard error of the mean). Representative photographs of pelleted WT and Δ*cydDC* after growth in hemin-containing medium (**E**). Intracellular heme concentration in *cydDC* genetic variants following growth in heme-containing medium (**F**, n=3, ** p < 0.01, ns = nonsignificant, ANOVA with Bonferroni correction).

This targeted validation of our positive selection screen yielded one transposon insertion mutant demonstrating significantly decreased intracellular hemin concentrations, relative to the other mutants and a WT control (tn_SAK_RS08800, **Fig. 3B**). This mutant had a transposon insertion in the first of a four-gene operon, *cydA-D*, annotated as a cytochrome oxidase system. The first two genes in the operon (*cydAB*) encode a terminal cytochrome oxidase that utilizes heme and menaquinone to drive respiratory metabolism ^18^, but the proteins do not have transmembrane domains or other surface-targeting features and therefore appeared unlikely to serve as heme importers.

However, the role of the *cydDC* half of the operon in GBS remained uncertain. A recent structural biology study described a CydDC ortholog in *E. coli* that transports heme embedded in the inner membrane to the periplasmic side where it is needed as a cofactor for cytochrome bd ^32^. The GBS CydDC heterodimer has a similar predicted structure as the *E. coli* system, with multiple transmembrane motifs. We therefore hypothesized that GBS *cydDC* encodes a heme transporter that potentially, as in *E. coli*, moves membrane embedded heme to the cytoplasmic space where it can be loaded into cytochrome bd, used for respiratory metabolism, dissociated from its central iron atom, and exported by HrtAB.

To validate our screen results, we generated a targeted deletion of *cydDC* in GBS (Δ*cydDC*) and a chromosomally complemented control (Δ*cydDC*::comp-*cydDC*) in which the gene had been replaced in-frame on the chromosome. We tested growth of these strains in inhibitory hemin (40 µM) concentrations, confirming that Δ*cydDC*, but not Δ*cydDC*::comp-*cydDC*, showed growth rescue consistent with the idea that deletion of a heme importer could lower intracellular oxidative stress under high-hemin conditions (**Fig. 3C-D**). During these experiments, we noted a phenotypic difference of Δ*cydDC*, which had an obvious lack of dark brown pigmentation compared to WT when grown in the presence of supplemental hemin, suggesting a difference in hemin sequestration between the two strains (**Fig. 3E**). As predicted by our hypothesis that CydDC is a heme importer, the Δ*cydDC* mutant showed decreased intracellular hemin concentrations relative to WT, an effect that could be *cis-*complemented by the full-length *cydDC* sequence (**Fig. 3F**).

### *CydDC* deletion causes intramembrane heme concentrations to increase

To test the hypothesis that CydDC encodes a heme importer that collects and transports heme to the intracellular space from an intramembrane pool, a function consistent with the recent structural-functional analysis of the *E. coli* ortholog ^32^, we performed direct measurement of heme from intracellular and membrane fractionation samples. Δ*cydDC*, but not the complemented control strain, showed increased intramembranous heme concentrations in supplemental hemin (**Fig. 4A**) and 20% lysed human erythrocytes (**Fig. 4B**). To confirm that *cydDC* and not *cydAB* was the necessary gene pair on the operon for importer activity, we also generated a Δ*cydAB* knockout and complemented it. The Δ*cydAB* strain showed modest growth and heme sequestration changes, each less than observed in Δ*cydDC* (**Suppl. Fig. 4**). A potential interpretation of this is that the CydAB cytochrome acts as the terminal heme receptor for the CydDC transporter, such that deletion of its locus leads to a secondary accumulation of heme due to blockade of CydDC heme release. Overall, these experiments are consistent with the CydDC protein pair that extracts heme for import from the cell membrane.

**Figure 4:**
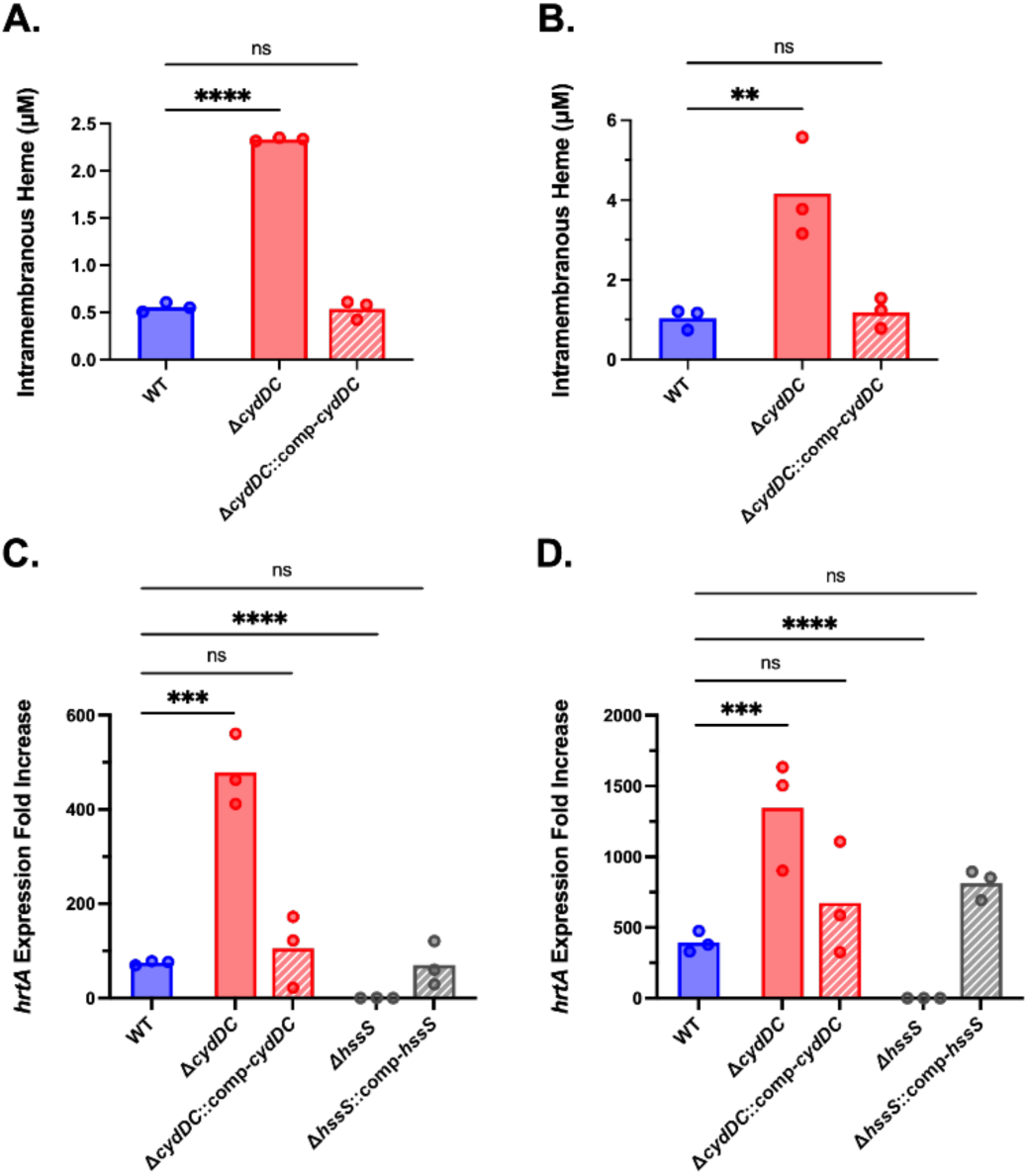
CydDC deletion leads to a buildup of intramembranous heme and overactivation of HssS signaling. Membrane fractionation and spectrophotometric detection of intramembrane heme was conducted in WT and *cydDC* genetic variants following exposure to hemin-containing medium (**A**) and lysed human erythrocytes (**B**). HssS signaling from heme binding to upregulate *hrtA* expression was measured in *cydDC* and *hssS* genetic variants following exposure to hemin-containing medium (**C**) and lysed human erythrocytes (**D**). n=3 for all conditions, ** p < 0.01, *** p < 0.005, *** p < 0.001, ns = nonsignificant, ANOVA with Bonferroni correction.

### HssRS signaling activity in Δ*cydDC* suggests intramembrane substrate sensing by HssS

Having determined through mutant testing that CydDC functions to extract heme from an intramembranous pool and deliver it to the intracellular compartment, we examined the influence of heme concentration modulation in those two cellular compartments on HssS signaling. Measuring *hrtA* expression by RT-qPCR as a readout of HssRS activity, we expected one of two possibilities. If the HssS heme binding site were localized intracellularly, the Δ*cydDC* mutant should show reduced signaling relative to WT and complement controls. However, if the HssS heme binding site were localized in the membrane space, accumulation of heme in the Δ*cydDC* membrane should increase HssRS-*hrtA* axis activity.

WT GBS and the Δ*cydDC* mutant were exposed to differing concentrations of exogenous hemin, and transcript levels of *hrtA* were quantified by RT-qPCR. Under basal conditions, WT and Δ*cydDC* strains showed comparable expression of *hrtA*. However, upon heme exposure, either as a purified additive (hemin) to TS medium (**Fig. 4C**) or as lysed human red blood cells (**Fig. 4D**), the Δ*cydDC* mutant exhibited significantly increased induction of *hrtA*, an effect that could be reversed by chromosomal complementation. To determine whether this augmented transcriptional response required HssS signaling, we performed analogous experiments in Δ*hssS* and complement control backgrounds. In the absence of HssS, heme-induced upregulation of *hrtA* was abolished, indicating that the enhanced signaling observed in the Δ*cydDC* mutant is strictly HssS dependent (**Fig. 4C-D**). Together, these data demonstrate that CydDC dampens HssS activation during heme exposure and support a model in which CydDC limits the availability of intramembranous heme accessible to HssS.

Additionally, we asked whether HssRS signaling reciprocally modulates expression of the *cydABCD* operon, which encodes the cytochrome bd oxidase (CydAB) and the CydDC heme importer. To address this, we quantified *cydA* transcript levels by RT-qPCR following exposure to either exogenous hemin or lysed human erythrocytes. In WT GBS, *cydA* expression was slightly induced under both conditions. This induction was attenuated in the Δ*hssS* mutant, suggesting partial dependence on HssRS signaling, whereas Δ*cydDC* showed equal or slightly increased *cydA* induction relative to WT (**Suppl. Fig. 5**). These effects were modest in magnitude compared to the robust HssRS-dependent regulation observed at the *hrtAB–hssRS* operon and were not sufficient to account for the pronounced phenotypes associated with loss of CydDC or HssS. Together, these data suggest that HssRS may exert limited transcriptional influence over the *cyd* operon under heme stress conditions but that regulation of *cydABCD* is likely dominated by additional inputs independent of canonical HssRS signaling.

### Structural and functional examination of GBS HssS reveal a novel heme binding site at the inner surface of the membrane lipid bilayer

To narrow the scope of potential heme binding sites on HssS, we performed protein-ligand interaction simulations with AlphaFold3 and Attracting Cavities 2.0 ^33,34^, which indicated presence of an inner membrane leaflet heme binding pocket formed at the intersection of the two transmembrane helices. Both modeling packages indicated protein-heme interaction sites at residues Y17, M21, H79, and R105 (**Fig. 5A-C**). We used a Cas12a-based mutagenesis system to generate chromosomally encoded alanine substitutions at these sites. After confirming the mutations by sequencing, we tested for HssS signaling activity by measuring hemin-inducible expression of the target *hrtAB-hssRS* operon. This experiment revealed greatest attenuation of HssRS signaling and hemin resistance demonstrated by the Y17A mutant, suggesting this residue as important for heme binding (**Fig. 5D**). Less dramatic, but still significant, reductions of hemin-induced *hrtA* expression were observed in the R015A and M21A mutants (**Fig. 5D**). As expected, the H149A control strain showed near-complete abrogation of hemin-inducible *hrtA* expression.

**Figure 5:**
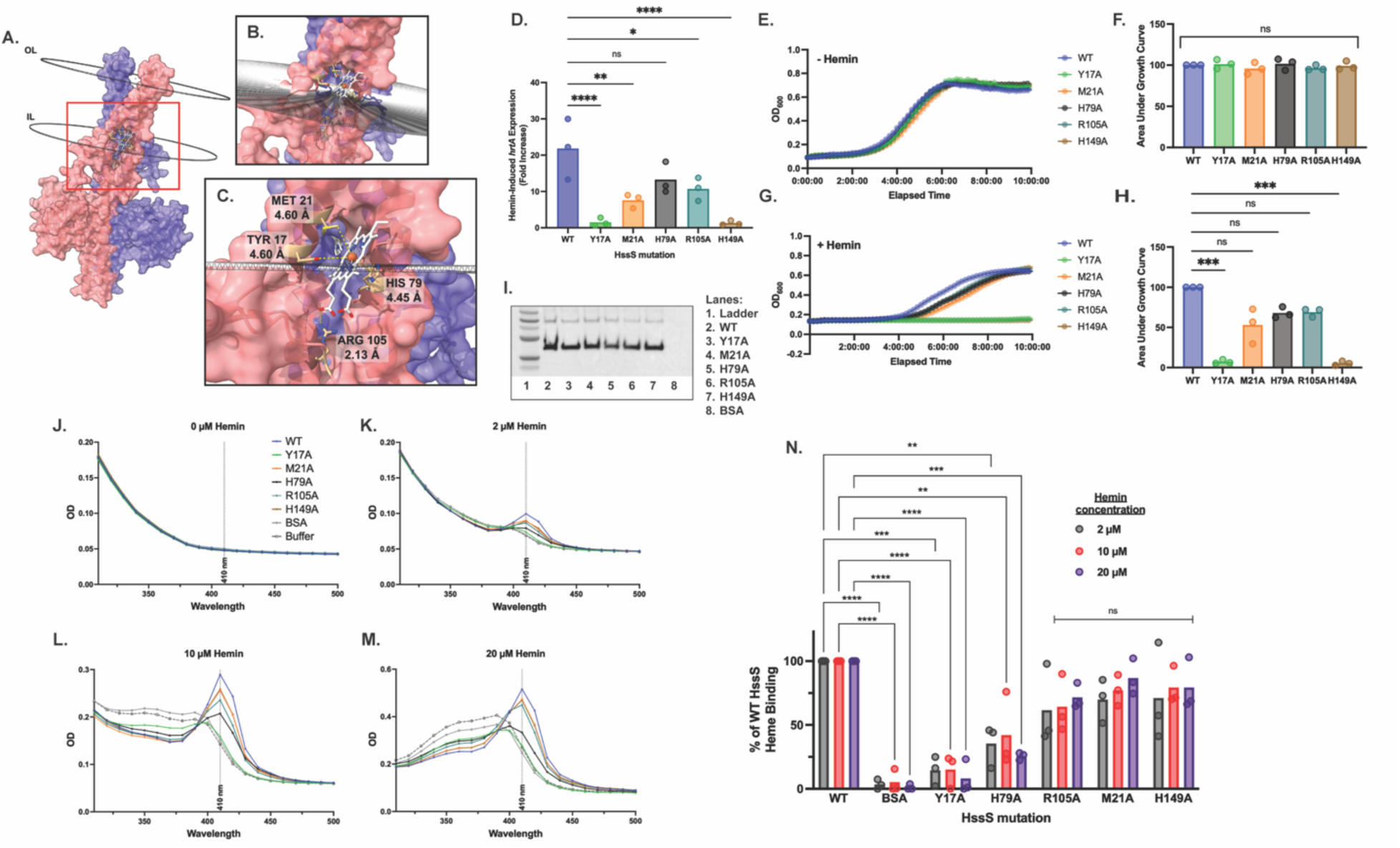
GBS HssS binds heme at an intramembranous pocket in the inner leaflet of the lipid bilayer. Docking models of heme interacting with the GBS HssS in its transmembrane configuration (OL = outer leaflet; IL = inner leaflet) show a putative binding pocket at the IMB (**A-B**). The nearest HssS amino acid residues, each estimated to stabilize heme at less than 5Å, are Met 21, Tyr 17, His 79, and Arg 105 (**C**). RT-qPCR of heme-induced HssS-mediated *hrtA* expression from chromosomally expressed targeted alanine substitution mutants of the four putative heme-stabilizing HssS amino acids, WT, and a H149A mutant that disrupts phosphorelay activity (**D**, n=3). Growth curves and areas under the curve of the chromosomally expressed HssS variants and WT in media without hemin (**E-F**, n=3) or with supplemental hemin (**G-H**, n=3). Statistical analyses for **F** and **H** were performed on raw AUC values and available in supplementals. We performed Polyhistidine-tagged, micelle-stabilized recombinant HssS binding site mutants were column affinity chromatography purified, diluted to equal concentrations, then visualized by Western blot (**I,** BSA = bovine serum albumin, negative control). Soret peak (410 nm, indicated) measurements were performed across a range of hemin concentrations (**J-M**). Areas under the Soret peak were normalized to WT and compared between mutants, WT, and BSA control. Statistical analyses were performed on raw OD_410_ readings available in supplementals (**N**, only significant comparisons shown). For all panels, * p < 0.05, ** p < 0.01, *** p < 0.005, **** p < 0.001, ns = nonsignificant, ANOVA with Bonferroni correction.

We also measured growth kinetics of our site mutagenesis strains in rich media and in hemin supplementation. Congruent with our *hrtA* RT-qPCR results, we found significant hemin-related growth attenuation in the Y17A experimental strain and the H149A positive control strain, with subtler growth defects in the remaining three targeted mutant strains (**Fig. 5E-H**).

The structural and functional studies described above suggest an inner layer intramembranous heme binding moiety in GBS HssS, which was consistent with findings from our domain swap mutant variants described above, where the strongest growth defects were seen from disruption of the inner domain and transmembrane domain 1, which together create the pocket defined by our *in silico* models. However, we remained cognizant that mutations altering coordination of the folded protein could disrupt heme responsive signaling and reduce GBS fitness in high heme conditions, thereby giving the illusion of a binding site. We therefore sought to establish a direct measure of heme binding to recombinant GBS HssS protein.

To directly assess heme binding by HssS, we measured Soret absorbance peaks, which provide a sensitive and specific spectral signature of heme coordination within protein environments ^35^. Free heme exhibits a characteristic absorbance maximum near 385 nm due to the π-π* transitions in the porphyrin ring system. Upon binding to proteins, this Soret band undergoes a red-shift and changes in amplitude, depending on the local environment and axial ligands presented by the protein. We purified recombinant WT and mutant HssS using detergent micelle solubilization, using Western blotting against the polyhistidine purification tag to confirm proper protein size and equivalent expression across strains (**Fig. 5I**). The Western blot, performed on protein preparations subjected to denaturing, reducing conditions, showed dominant bands of the proper size for HssS monomers, as well as a larger molecular weight band that faded with harsher denaturing conditions, suggesting that it reflected dimerized HssS (**Suppl. Fig. 6**).

We reconstituted the purified protein micelles with equimolar hemin, using WT as a positive control and bovine serum albumin (BSA, which can engage in nonspecific hydrophobic heme interactions but does not have a specific heme-binding site) as a negative control. UV-Visible spectroscopy revealed a red-shift of the Soret band to ∼410 nm upon incubation with WT HssS (**Fig. 5J-M**), consistent with stable coordination of heme by the protein. No 410 nm peak was observed in buffer+hemin or BSA+hemin control conditions, in which only the characteristic spectrum of dissolved hemin was measured (**Fig. 5J-M**).

In contrast to WT, alanine substitution mutants at the predicted binding site residues Y17 and H79, found within transmembrane domains 1 and 2, respectively, displayed significantly reduced peak shift and diminished amplitude, suggesting impaired or absent heme binding. These spectral changes were quantified by difference spectroscopy (**Fig. 5J-M**), and the relative heme-binding capacities of the mutants were further confirmed by plotting the absorbance at 410 nm against protein concentration (**Fig. 5N**). In aggregate, these findings support a model in which specific intramembranous residues, especially Y17, contribute to the formation of a heme-binding pocket within the HssS transmembrane region. Protein sequence comparison to HssS in closely related bacterial relatives demonstrates that the Y17 and H79 are modestly conserved (**Suppl. Fig. 7**), suggesting a shared role in functional heme interactions.

### A mouse sepsis model demonstrates importance of the HssRS and CydDC systems in virulence

The *in vitro* experiments described above indicated that coordinated sensing and import of heme at the cell membrane are critical for maintaining intracellular heme homeostasis during exposure to blood-associated heme. To determine whether disruption of this intramembranous heme regulatory system compromises GBS fitness in physiologically relevant settings, we next examined the contributions of HssS and CydDC to virulence *in vivo*.

WT Balb/C mice were infected by tail vein injection with equal inoculates of WT, Δ*hssS*, and Δ*cydDC* GBS strains and monitored for survival, weight changes, and bacterial burden. WT GBS developed progressive illness and high-grade bacteremia, whereas mice infected with either Δ*hssS* or Δ*cydDC* exhibited complete survival (**Fig. 6A**) and weight retention (**Fig. 6B**). Consistent with these outcomes, bacterial burdens recovered from blood and highly perfused organs—including liver, kidney, and brain—were markedly reduced in mice infected with either mutant strain compared to WT (**Fig. 6C-J**).

**Figure 6:**
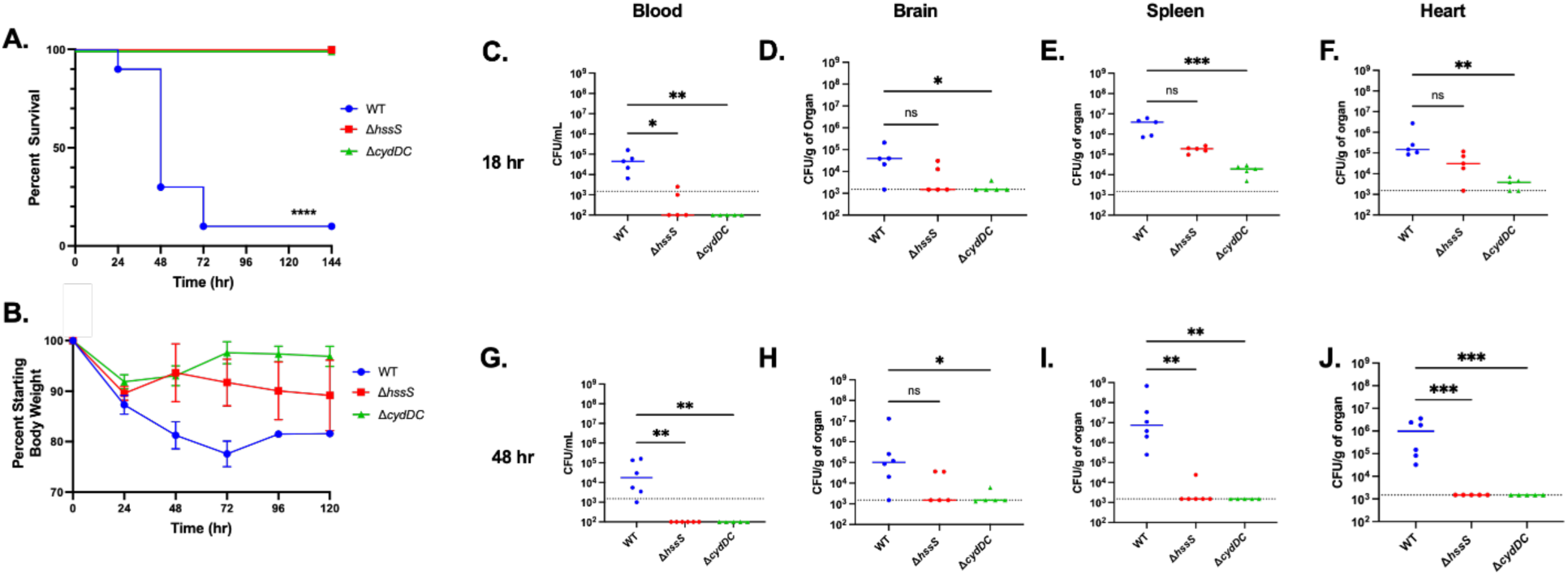
Both HssS and CydDC are required for virulence in a mouse bacteremia model. In a model of bacteremia, adult Balb/C mice (n=10) were intravenously administered 10^7^ WT, Δ*hssS*, or Δ*cydDC* and monitored for survival and weight loss over a six-day period (**A-B**, **** p < 0.001 Mantel-Cox). In a separate experiment, infected mice were euthanized at 18 and 48 hours and tissues harvested for GBS CFU quantification (n = 5 for mutants, n = 4 or 6 in WT, reflecting variable survival among an initial n = 10, **C-J**, * p < 0.05, ** p < 0.01, *** p < 0.005, ns = nonsignificant, Kruskal-Wallis with Dunn’s correction).

Importantly, the virulence attenuation observed for Δ*hssS* and Δ*cydDC* was comparable in magnitude, despite the distinct molecular roles of these proteins in heme sensing and import, respectively. These findings support a model in which both functions are essential components of a unified heme homeostasis pathway required for intravascular persistence. Together, the *ex vivo* and *in vivo* data demonstrate that disruption of either heme sensing at the membrane (HssS) or extraction of membrane-associated heme for intracellular use (CydDC) compromises GBS survival in blood and attenuates systemic infection.

### Model proposal: a membrane heme pool serves as a thermostatic reservoir for HssS sensing and CydDC import

The collective findings from our genetic, biochemical, and transcriptional analyses support a model in which heme partitions into the GBS cell membrane, forming a dynamic reservoir that integrates sensing, import, and export. In this framework, intramembranous heme serves as the immediate substrate for both HssS-dependent signaling and CydDC-mediated import, with HssRS-driven export, through HrtAB, acting to prevent toxic accumulation. We therefore propose that the membrane heme pool functions as a “thermostatic” regulator, buffering fluctuations in environmental heme availability and coordinating downstream responses (**Fig. 7**).

**Figure 7:**
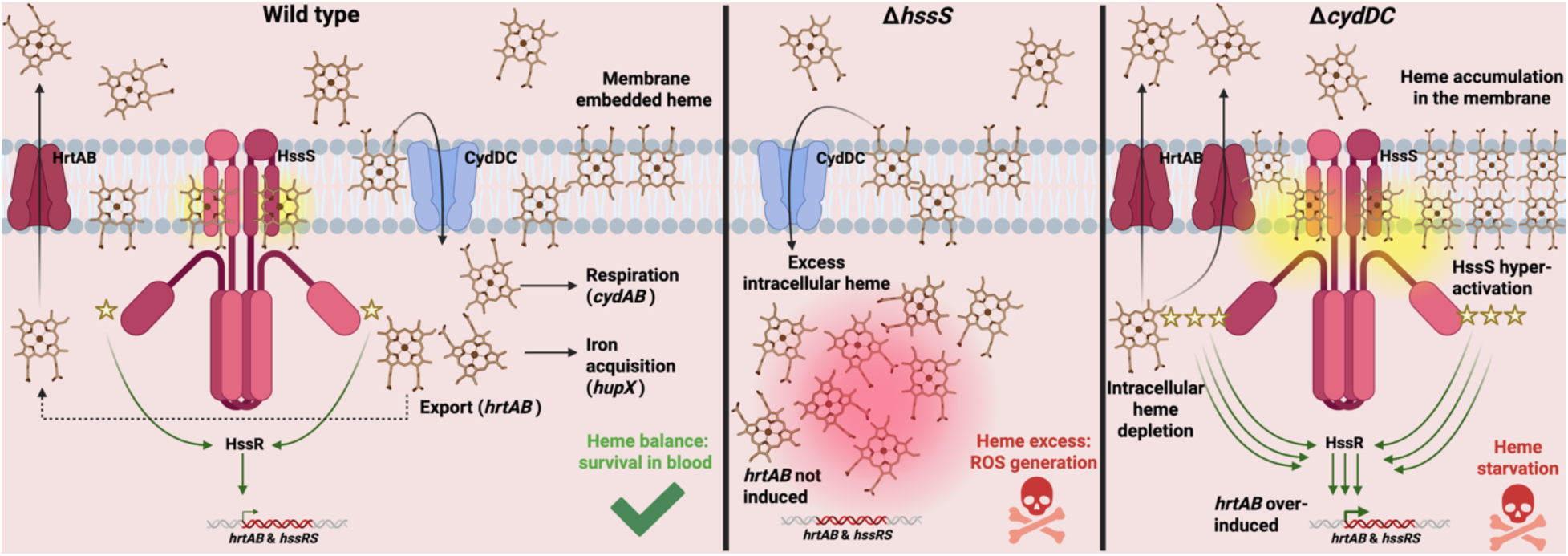
Proposed model of membrane-based heme storage and utilization for GBS homeostasis during bacteremia. In this conceptualization, WT GBS (left panel) uses its cell membrane lipid bilayer as a thermostatic reservoir for heme storage and sensing; CydDC continuously imports membrane-embedded heme into the intracellular space for use in respiration and iron acquisition, while the HssS intramembrane heme binding pocket senses and responds to heme buildup by auto-upregulating and increasing expression of the HrtAB heme export system. Deletion of *hssS* (middle panel) creates a toxic buildup of heme due to failure to counteract CydDC importation through a balanced upretulation of the HrtAB exporter. Deletion of *cydDC* (right panel) causes overaccumulation of heme in the membrane, leading to hyperactivation of HssS signaling, further driving down intracellular heme levels and creating lethal shortage of substrate for heme-dependent biochemistry (**C**).

In WT GBS, heme derived from the extracellular environment equilibrates into the lipid bilayer, where it is available for detection by the intramembranous HssS sensor and extraction by the CydDC transporter. These processes operate in concert: CydDC mobilizes heme from the membrane for intracellular liberation of iron by a heme oxygenase, HupZ ^31^, and coordination to CydAB for respiration, while HssS monitors membrane heme abundance and activates the HrtAB export system when local concentrations rise above a tolerable threshold. In this way, WT GBS maintains a balanced distribution of heme across membrane and intracellular compartments, permitting metabolic utilization while avoiding toxicity.

Disruption of either arm of this system is predicted to destabilize heme balance in distinct ways. In the absence of HssS signaling (**Fig. 7**), impaired induction of HrtAB-mediated export results in inappropriate retention of heme and increased susceptibility to heme-mediated stress. Conversely, loss of CydDC (**Fig. 7**) leads to heme accumulation within the membrane, leading to exaggerated HssS activation and reduced intracellular heme availability. These perturbations predict differential fitness consequences depending on environmental heme availability.

To test this model, we examined growth and survival of WT and mutant strains in *ex vivo* human blood lysate (**Fig. 8A-D**) and whole blood (**Fig. 8E**).

**Figure 8:**
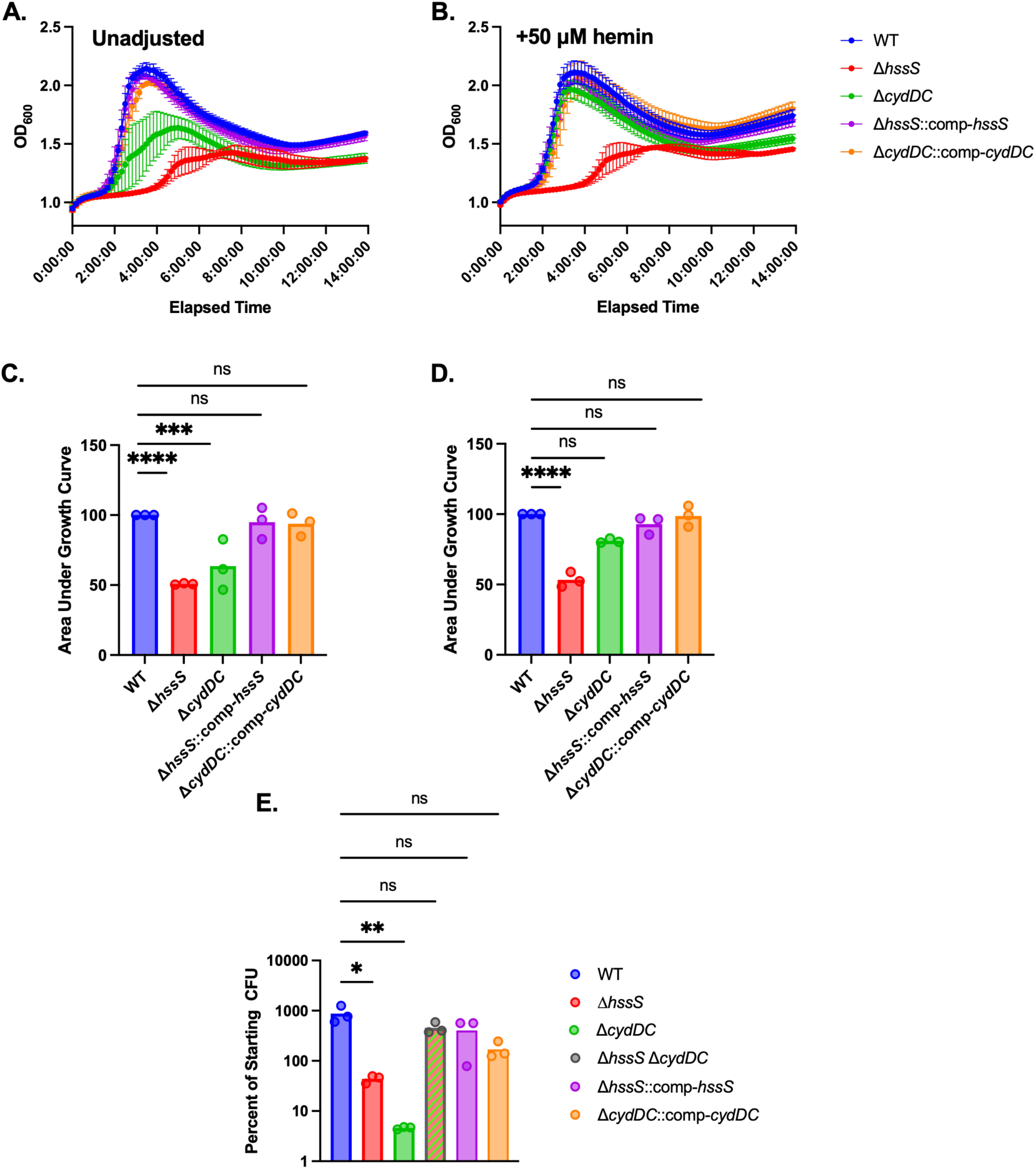
Model testing in human blood and lysates with variable heme concentrations and GBS genotypes. Genetic *hssS* and *cydDC* GBS mutants and their complements were grown in a clarified lysate of human erythrocytes (**A**) and clarified lysate supplemented with 50 μM hemin (**B**) and quantified by area under the growth curves (**C-D** unsupplemented and hemin supplemented, respectively; n=3). Survival of single mutants, complemented single mutants, or the double mutant Δ*hssS* Δ*cydDC* were tested after 6-hour incubation in fresh human whole blood (**E**). n=3, * p < 0.05, ** p < 0.01, *** p < 0.005, **** p < 0.001, ns = nonsignificant, error bars denote standard error of the mean, ANOVA with Bonferroni correction.

In human whole blood lysate, which is expected to contain relatively low concentrations of freely available heme due sequestration mediated by hemopexin and hemoglobin, both Δ*hssS* and Δ*cydDC* mutants exhibited impaired growth compared to WT, as assessed by growth kinetics (**Fig. 8A-B**) and area under the growth curve (**Fig. 8C-D**). These findings are consistent with the model prediction that, under heme-limited conditions, both efficient sensing/export (HssS) and membrane extraction/import (CydDC) are required to maintain sufficient intracellular heme for metabolic needs.

We next asked whether increasing extracellular heme availability could bypass the requirement for membrane extraction by CydDC. Supplementation of blood lysate with 50 μM hemin restored growth of the Δ*cydDC* mutant to near-WT levels (**Fig. 8B, D**), supporting the interpretation that high ambient hemin concentrations enable sufficient passive partitioning and/or alternative uptake mechanisms to compensate for loss of CydDC function. In contrast, Δ*hssS* mutants remained impaired under these conditions, consistent with a persistent defect in heme-responsive regulation and detoxification that cannot be overcome (and may be worsened) by increased substrate availability. Additionally, we tested Δ*cydAB* growth in the two experimental conditions described above (**Suppl. Fig. 8**). We found that, in contrast to Δ*cydDC*, the cytochrome mutant grew comparably to WT in whole blood lysate but had a growth defect in whole blood lysate supplemented with 50 μM hemin. This indicates distinct and complementary roles for the two systems, CydDC and CydAB, with the cytochrome CydAB potentially playing a detoxifying role in settings of elevated heme concentrations in the surrounding hematologic environment.

Finally, to evaluate how these pathways interact under physiologic conditions, we assessed bacterial survival in freshly collected human whole blood. As observed previously, both Δ*hssS* and Δ*cydDC* mutants demonstrated significantly reduced survival compared to WT (**Fig. 8E**), rescuable by complementation, reflecting the importance of coordinated heme sensing and import during bloodstream exposure. Strikingly, however, a double knockout strain (Δ*hssS* Δ*cydDC*) lacking both HssS and CydDC showed a partial restoration of survival relative to either single mutant.

This result is consistent with a model in which loss of CydDC-mediated import reduces intracellular heme burden, partially offsetting the toxicity that would otherwise result from impaired HssS-dependent export. Conversely, absence of HssS signaling may limit induction of heme export pathways that would otherwise deplete intracellular heme in the setting of reduced import. Thus, simultaneous disruption of both pathways may rebalance heme distribution across compartments, restoring a degree of functional homeostasis despite loss of regulatory control.

Taken together, these findings provide experimental support for a model in which the GBS membrane serves as a central hub for heme homeostasis. By acting as a reservoir that buffers environmental fluctuations and coordinates sensing, import, and export, the membrane heme pool enables GBS to adapt to the complex and dynamic conditions encountered during bloodstream infection.

## Discussion

In this study, we identify the cytoplasmic membrane as a site of heme accumulation, sensing, and redistribution in GBS during bloodstream invasion. While many prior models of bacterial heme homeostasis have treated heme as a molecule that is either extracellular, requiring active capture and transport, or cytosolic, where it is incorporated into metabolism or exported to limit toxicity, our findings support a third, intermediate state in which heme partitions into the membrane lipid bilayer and serves as a functional signaling and metabolic reservoir. We show that GBS exploits this intramembranous heme pool through two coordinated processes: sensing via the membrane-embedded histidine kinase HssS, and extraction for intracellular use via the CydDC transporter. Disruption of either process compromises survival in blood and attenuates systemic infection, indicating that membrane-centered heme regulation is a critical determinant of intravascular fitness.

Our data further establish HssS as an intramembranous heme sensor that detects heme accumulation within the membrane rather than relying on extracellular ligand binding. Unlike previously characterized HssS orthologs in *Staphylococcus aureus* and *Bacillus anthracis*, which fold substantial extracellular domains, GBS HssS lacks a robust extracellular domain to participate in heme stabilization and binding. Through a combination of structural modeling, targeted mutagenesis, and biochemical assays, we identify specific inner membrane leaflet-associated residues required for heme binding and signaling and demonstrate heme interactions by purified recombinant HssS protein. These findings suggest that heme insertion into the membrane constitutes the relevant trigger for downstream HssR signal transduction.

In parallel, we identify CydDC as a membrane-associated heme extraction system that limits intramembranous heme availability and promotes intracellular heme utilization. Loss of CydDC leads to accumulation of heme in the membrane, heightened HssS signaling, and reduced intracellular heme levels, consistent with a role in mobilizing membrane-embedded heme for downstream pathways. While CydAB has been recognized as encoding a heme-dependent terminal oxidase that supports respiratory metabolism in GBS, the function of CydDC has remained unclear. Our findings place CydDC within a broader heme homeostasis circuit in which membrane-associated heme is dynamically balanced between sensing, metabolic use, and detoxification. Together, these observations support a model in which intramembranous heme serves as a shared substrate for both regulatory and metabolic processes during bloodstream exposure, enabling GBS to exploit host-derived heme without requiring dedicated surface capture systems.

This model is consistent with the amphipathic properties of heme, which favor spontaneous partitioning into lipid environments. While membrane association of porphyrins has been described in other systems ^32,36,37^, the concept has not been deeply explored in the context of bacterial pathogen homeostasis. A pioneering recent study in this regard showed that *S. aureus* HssS contains key arginine and phenylalanine residues that stabilize binding in a pocket at the interface between the extracellular domain and the outer surface of the cell membrane lipid bilayer ^24^. In contrast, GBS HssS appears to sense heme abundance deeper in the lipid bilayer through a binding pocket embedded at the inner layer of the cell membrane. Such a mechanism may be particularly advantageous in environments like the bloodstream, where fluctuations in free heme concentration may be rapid and spatially heterogeneous, but where membrane partitioning could provide a more stable and integrative readout of exposure.

Our results suggest that, in organisms lacking dedicated extracellular heme capture systems, passive equilibration into the membrane may serve as a primary route of acquisition, with subsequent extraction and sensing occurring directly within the bilayer. This mechanism provides an efficient means of exploiting host-derived heme without reliance on specialized surface receptors, which are absent in GBS and variably distributed across related gram-positive species. Our findings open potential for examining whether heme reservoir sequestration in the bacterial membrane is a mechanism used by other species associated with bacteremia but lacking obvious genetic pathways for heme acquisition.

An important implication of the membrane-HssS-CydDC axis is that heme homeostasis in GBS is governed not solely by absolute heme abundance but by its distribution across cellular compartments. This is highlighted by the differential phenotypes observed in single and double mutants of *hssS* and *cydDC*. Whereas loss of either gene alone disrupts survival in blood, simultaneous disruption of both pathways partially restores fitness, suggesting that rebalancing of heme pools can compensate for loss of regulatory control. These findings underscore the importance of considering intracellular partitioning, rather than environmental concentration, as the key determinant of heme toxicity and utility.

GBS-mediated hemolysis has been the focus of extensive study. The cytolytic ornithine-rhamnopolyene molecule β-hemolysin/cytolysin ^38–40^, whose biosynthetic *cyl* operon is tightly coordinated by the master virulence regulatory CovRS system ^41–46^, is known to have a significant role in determining GBS virulence across numerous anatomic settings, including during bacteremia. β-hemolysin/cytolysin is believed to intercalate into eukaryotic lipid bilayers, causing osmotic lysis and releasing intracellular contents ^47^. Coincubation of GBS with human erythrocytes causes hemolysis and release of free hemoglobin in a β-hemolysin/cytolysin-dependent fashion ^39,41,45,47^. While most erythrocyte intracellular heme is bound to hemoglobin and free heme is quickly sequestered in the bloodstream by hemoproteins, we consider it feasible that local release of free heme at the site of erythrocyte breakdown may be enough to “charge” the GBS cell membrane, establishing a portable, protected heme source for GBS to utilize as it spreads systemically. One potential topic of future study, beyond the scope of this project, would be to examine heme capture by GBS in the immediate vicinity of β-hemolysin/cytolysin-mediated hemolysis.

While our findings define a membrane-centered mechanism for heme homeostasis in GBS, several areas warrant further investigation. Our biochemical fractionation and functional analyses consistently support accumulation of heme within the membrane, establishing this compartment as a key site of heme availability; future studies employing higher-resolution approaches may further refine the spatial organization and dynamics of this pool in situ. In addition, our data identify CydDC as a principal mediator of heme extraction from the membrane for intracellular use, though it is likely that downstream trafficking involves additional components that remain to be defined. Finally, although disruption of this system markedly attenuates virulence in a murine model of bacteremia, the precise host-derived sources and temporal dynamics of heme exposure during infection represent an important area for future study.

In summary, we propose that intramembranous heme partitioning represents a central organizing principle for heme homeostasis in GBS during bloodstream infection, a concept that may extend to other blood borne bacteria that lack dedicated extracellular heme acquisition pathways. By coupling membrane-based sensing, extraction, and export, this system enables bacteria to buffer fluctuations in environmental heme availability while maintaining access to this essential but potentially toxic cofactor. These findings expand current models of bacterial heme biology and suggest that the membrane may serve as a critical intermediate compartment for the regulation of other hydrophobic metabolites encountered during host-pathogen interactions.

## Methods and Materials

### Ethics Statement

Animal experiments were performed under an approved IACUC protocol at University of Pittsburgh. Phlebotomy for hemolysis assays was conducted under a University of Pittsburgh approved IRB protocol following obtaining informed consent from healthy adult volunteers.

### Bacterial Strains, Plasmids, and Growth Conditions

GBS serotype Ia strain A909 was used as the WT background. Bacteria were routinely cultured in tryptic soy broth (TSB) at 37 °C under static conditions. Where indicated, cultures were supplemented with hemin at concentrations specified.

All plasmids were maintained using erythromycin (Erm) selection (5 µg/mL GBS and 300 µg/mL *E. coli*^)^. Clean deletion mutants (Δ*hssS*, Δ*hssRS,* Δ*cydDC,* Δ*cydAB*, and Δ*hssRS*Δ*cydDC*) were generated using a published Cas12a-mediated genome editing strategy ^48^. Chromosomal *hssS* mutant alleles were generated using the same Cas12a-based approach. Complementation was performed by Cas12a-mediated restoration of the deleted locus (in *cis*) at its native chromosomal position, reverting the knockout allele to the WT sequence. Markerless editing was confirmed by PCR and sequencing.

For recombinant protein expression, an *E. coli* T7 expression strain (NEB C3013I) was cultured in Luria–Bertani (LB) medium with kanamycin (Kan) selection (50 µg/mL) and induced as described below.

### Domain Mutants

Transmembrane domain boundaries of HssS were defined using the DeepTMHMM server ^49^. Domain swap mutans, in which the transmembrane domain 1 or 2 of GBS HssS was replaced with a duplication of the other transmembrane, were designed based on these predicted boundaries. Alanine block substitution mutants were generated by replacing residues of interest with alanine codons; to minimize sequence repetitiveness and reduce the likelihood of recombination artifacts, all four synonymous alanine codons (GCT, GCC, GCA, GCG) were used in random combination across each substitution block. All mutant *hssS* alleles were ordered as synthetic gene fragments and cloned into the pGBScomp complementation vector by Gibson assembly. The resulting plasmids were introduced into a Δ*hssS* background and maintained under erythromycin selection.

### Preparation of Hemin Stocks

Hemin stocks were prepared fresh for each experiment by dissolving 97% porcine hemin chloride (Thermo #A11165.06) in DMSO to generate 10 mM stocks, which were subsequently diluted as required. All hemin-containing solutions were protected from light throughout preparation and use. Hemin refers specifically to the oxidized form of the molecule (ferric) that is easily obtainable commercially while heme refers to the reduced form of the molecule.

### CRISPR Interference (CRISPRi) Negative-Selection Screen

A curated CRISPRi knockdown library targeting two-component system (TCS) operons in *S. agalactiae* A909 expressing catalytically inactive Cas9 (A909 *dcas9*) was used for negative-selection screening ^27^. Each TCS was targeted with two independent sgRNAs designed using the Broad Institutes CRISPick ^50^ to design protospacer targets. Each sgRNA was inserted into the p3015b sgRNA expression plasmid and electroporated into A909 *dcas9* strain as an individual CRISPRi strain. These individual CRISPRi strains were equally pooled in the CRISPRi knockdown library by resuspending colonies of individual knockdown strains to an OD_600_ of 1 in PBS and adding 100 µL of each strain together.

An aliquot of the pooled library was collected as the input sample, after which parallel aliquots of the library were exposed either to 10 µM hemin or to unsupplemented TSB as a no-heme control to account for general fitness effects. Following exposure, surviving bacteria were collected as output samples. Plasmid DNA was isolated from input and output samples, and the sgRNA protospacer region was amplified for 22 cycles by PCR and subjected to amplicon sequencing on an Illumina MiSeq platform.

Sequencing reads were normalized to total reads within each biological replicate to generate population percentages for each protospacer within that replicate. Fold changes were calculated by dividing the output population percentage by the input population percentage for each protospacer within each condition. To isolate hemin-specific fitness effects, the fold change observed under hemin exposure was subtracted from the fold change observed in the no-hemin control for each protospacer, yielding a Δfold change value where negative values indicate reduced strain abundance under hemin stress and positive values indicate relative enrichment. Protospacer values were averaged across the two independent sgRNA expressing strains targeting each TCS operon, and results represent the mean of three independent biological replicates.

PCA analysis was performed in Python using the Scikit-Learn library. Raw population percentages were averaged across protospacers and standardized by z-score normalization using scikit-learn StandardScaler. PCAs were performed using the Scikit-Learn’s PCA function and the first two principal components were retained for visualization. Figures were generated using matplotlib. Annotated python scripts are available upon request.

### Transposon Library Positive-Selection Screen

A previously described, saturated GBS transposon mutant library was subjected to positive selection under lethal hemin stress conditions ^28^. The pooled library was exposed to 100 µM hemin, and surviving colonies were isolated. Transposon insertion sites in the surviving variants were identified by targeted sequencing of genomic DNA using a transposon-specific initiation primer (TN917 faces US, **Suppl. Data 1**). Candidate genes disrupted by transposon insertion were evaluated for predicted transporter function and membrane localization.

### Intracellular Heme Quantification

Intracellular heme concentrations were measured using an oxalic acid–based fluorescence assay ^51^. The following steps were protected from light as much as possible. Briefly, bacterial cultures were grown overnight in TSB with or without 10 µM hemin, pelleted by centrifugation, and washed twice in phosphate-buffered saline (PBS). Cell pellets were resuspended in lysis buffer containing 10mM Tris-HCl (pH 7.5), 250U/mL mutanolysin, 2 mg/mL lysozyme, 5 µL/mL DNase I, and 5 mM MgCl₂ and incubated at 37 °C for 1-hour gentle rocking.

Cells were mechanically lysed by bead beating using 0.1 mm zirconia/silica beads, followed by clarification at 10,000 × g. Supernatants were subjected to ultracentrifugation at 100,000 × g to separate cytosolic and membrane fractions. Total protein concentrations were determined using a Bradford assay and used as a total lysis control. For heme quantification, lysates were mixed 1:1 with 2 M oxalic acid. Samples were split into boiled and unboiled aliquots, boiled for 5 minutes where indicated, and transferred to black clear-bottom plates. Unboiled samples were used for background normalization. Fluorescence was measured using excitation at 400 nm and emission at 608 nm, with spectral scans collected from 400–700 nm. Heme concentrations were interpolated from standard curves and normalized to total protein content (Bradford Assay: Thermo #23200).

### Membrane Fractionation and Membrane-Associated Heme Measurement

For membrane-associated heme measurements, mid-logarithmic growth cultures (OD₆₀₀ ≈ 0.6) were harvested and washed 3 times in 1x Tris-HCl buffer (pH 7.5). Cell walls were digested enzymatically (same lysis buffer as intracellular heme measurements, see above) followed by mechanical lysis via bead beating. Clarified lysate supernatants were subjected to ultracentrifugation at 100,000 × g for 2 hours at 4 °C to pellet membrane fractions.

Membrane pellets were resuspended in 1x Tris-HCl buffer, and heme content was quantified using the same oxalic acid fluorescence assay as described above. Bradford assays were performed on resuspended membrane pellets. Heme concentrations were interpolated from standard curves and normalized to total protein content (Bradford Assay: Thermo #23246).

### RNA Isolation and Quantitative RT-PCR

Total RNA was extracted using a NIMBUS Presto automated liquid handling system employing a modified MagMax-based protocol (Fischer: #A42356) optimized for RNA isolation from GBS. GBS optimization of this protocol includes the addition of mutanolysin during enzymatic lysis and a 1:10 concentration of DNA binding beads in the bead binding buffer. Immediately following extraction, RNA samples were treated with DNase (2 µL DNase per 40 µL RNA) at 37 °C for 2 h, followed by heat inactivation 75 °C for 10 minutes. cDNA synthesis was performed using standard reverse transcription kit (Fisher: #4368814). Quantitative PCR was carried out using gene-specific primers, with transcript levels normalized to the housekeeping gene *srtA* (Sortase A). Relative gene expression was calculated using the ΔΔCt method and expressed as fold-change relative to untreated control conditions.

### Recombinant Expression and Purification of HssS

Our method was adapted from a standardized histidine kinase purification protocol ^52^. Full-length HssS and variant proteins were expressed in *E. coli* T7 expression strains harboring His₆-tagged constructs cloned into the pET28a expression vector. 1 L cultures were grown with shaking in baffled flasks at 37 °C to an optical density at OD₆₀₀ of 0.1 to 0.2 and induced with 1 mM isopropyl β-D-1-thiogalactopyranoside (IPTG) and 2 mM lactose. Following induction, incubation temperatures were dropped to 25 °C and cultures were allowed to grow overnight to promote membrane protein expression with reduced toxicity and reduced aggregation.

All steps performed from here were kept chilled at 4 °C. Cells were harvested by centrifugation, resuspended in chilled lysis buffer (50 mM Tris-HCl [pH 8], 300 mM NaCl, 10% glycerol), and disrupted by bead beating using 0.1 mm zirconia/silica beads. Bead beating steps were performed at 4 °C. Cell debris was removed by low-speed centrifugation, and membranes were isolated by ultracentrifugation of supernatants at 100,000 × g for 2 hours at 4 °C. Membrane pellets were solubilized in chilled solubilization buffer containing 50 mM Tris-HCl (pH 8), 300 mM NaCl, 30% glycerol, 20 mM n-dodecyl β-D-maltoside (DDM), 20 mM imidazole and solubilized proteins were purified by nickel affinity chromatography (Qiagen #30410). Bound proteins were eluted in 8 fractions (200 µL at a time) using elution buffer (50 mM Tris-HCl [pH 8], 300 mM NaCl, 30% glycerol, 4 mM DDM, 300 mM imidazole) and analyzed by SDS–PAGE. Fractions containing the highest yield and purity were pooled and serially dialyzed against dialysis buffer containing 50 mM HEPES buffer, 50% (vol/vol) glycerol and 1mM DDM. Purified proteins were stored for no longer than 24 hours at 4°C, and protein concentrations were determined spectrophotometrically using calculated extinction coefficient (GBS HssS: 37,360 M⁻¹·cm⁻¹).

### Soret Shift Heme-Binding Assays

Purified HssS proteins were diluted to 2 µM in dialysis buffer and incubated with increasing concentrations of hemin prepared in matched buffer. Reactions were assembled in microplate format and gently mixed prior to measurement. Absorbance spectra were collected from 300–700 nm using a plate reader, and Soret shifts were monitored as an indicator of heme binding. Dialysis buffer and 2 µM bovine serum albumin (BSA) suspended in dialysisbuffer served as controls.

### SDS–PAGE and Western Blotting

Purified proteins were separated by SDS–PAGE using Bis–Tris gels (Fisher NW04120BOX) and transferred to polyvinylidene difluoride (PVDF) membranes. Membranes were stained with Ponceau S to confirm transfer, rinsed with water, blocked, and probed with anti-His primary antibodies followed by horseradish peroxidase–conjugated secondary reagents (Thermo #35061). Signals were detected by chemiluminescence and imaged using a ChemiDoc imaging system.

### Human Whole Blood Survival Assays

Fresh human whole blood was obtained from healthy adult volunteers under an IRB–approved protocol and anticoagulated with heparin sulfate. Bacterial strains were OD₆₀₀ normalized to 1 in 1x PBS before 100 µL inoculation into 1mL of blood and incubated at 37 °C on a rocking platform to maintain gentle mixing. At designated time points, samples were serially diluted and plated to count surviving CFU.

### Mouse Infection Experiments

Female BALB/c mice aged 8–10 weeks were used for all in vivo experiments performed under an approved IACUC protocol. Mice were infected via intravenous tail vein injection with 1 × 10⁷ CFU of the indicated GBS strains. Disease progression was monitored by survival analysis, and bacterial burdens were quantified in organs harvested 18 and 48 hours post-infection.

### Statistical Analysis

All experiments were performed with a minimum of three independent biological replicates. *In vitro* studies used a minimum two technical replicates of measurements, for which the average was used as the reported value of the independent biological replicate. Where indicated, biological replicates represent the mean of three technical replicates. Statistical analyses were performed GraphPad Prism. Exact statistical tests and sample sizes are indicated in figure legends.

### Protein Modeling and Ligand Binding

Structural models of HssS were generated using AlphaFold3 via the AlphaFold Server with default multiple sequence alignment settings and five model seeds ^34^. The top-ranked model was selected based on predicted local distance difference test (pLDDT) scores and used for all downstream analyses. Transmembrane topology was predicted using the Deep TMHMM server and cross-referenced with the AlphaFold3 structural model to annotate domain boundaries, including the two transmembrane helices, the short extracellular bridge, the intracellular N-terminus, and the cytoplasmic HAMP and DHp/CA signaling domains. Molecular visualization and structural figures were prepared using UCSF ChimeraX.

For *in silico* ligand docking, the heme digital model was obtained from the RCSB PDB Chemical Component Dictionary and docked into the putative intramembranous binding region of the HssS structural model using Attracting Cavities 2 through the SwissDoc server ^33,53^. AlphaFold3 was also used to predict heme docking using heme in the preloaded ligand library of the AlphaFold Server. Predicted heme interacting residues were consistent across both models and within five angstroms of heme.

### Clarified Lysed Whole Blood Media

Heparinized human whole blood was centrifuged to separate plasma and cellular fractions. Plasma was removed and retained at 4 °C or −80 °C, and packed red blood cells (RBCs) were collected. RBCs were osmotically lysed by dilution 1:1 with sterile water and incubated for 30 minutes at room temperature or overnight at 4 °C. Lysed RBCs were combined with serum to a final composition of 60% lysed RBC fraction and 40% serum and incubated for 30 minutes at room temperature with gentle rocking. The mixture was clarified by centrifugation at 10,000 × g for 5 minutes, and the supernatant was collected. The supernatant was passed through a 0.22 µm filter and the resulting solution was used as clarified lysed whole blood media.

### Growth Curves

Bacterial strains were OD₆₀₀ normalized to 1 in 1x PBS before 1:100 inoculation.Cultures were incubated at 37 °C and OD₆₀₀ was recorded at regular intervals using a plate reader. Growth curves were performed in technical triplicate within each biological replicate. Area under the curve (AUC) values were calculated using the trapezoidal method as a summary metric of overall growth. All statistical analyses were performed on raw AUC values before normalization to control strain.

For growth in clarified whole blood, cultures were inoculated 1:100 into clarified whole blood medium prepared as described above and monitored under identical conditions. Where indicated, clarified whole blood was used alone or supplemented with 50 µm hemin as specified in figure legends. Media-only controls were included for each condition and used for background subtraction.

## Acknowledgements

Dr. Laura Mike generously provided valuable advice on HssS purification and heme detection assays. We thank Dr. Devin Stauff for providing insights on HssS heme binding and for providing access to a plate reader spectrophotometer. We are grateful to Drs. Terrence Dermody and Danica Sutherland for facilitating use of an ultracentrifuge. Dr. William Macdonald and the University of Pittsburgh Health Sciences Sequencing Core provided excellent assistance and troubleshooting with CRISPRi library sequencing. This research was supported in part by the University of Pittsburgh Center for Research Computing and Data, RRID:SCR_022735, through the resources provided. Specifically, this work used the HTC cluster, which is supported by NIH award number S10OD028483. Figures 2B and 7 were created in BioRender.com.

## Funding

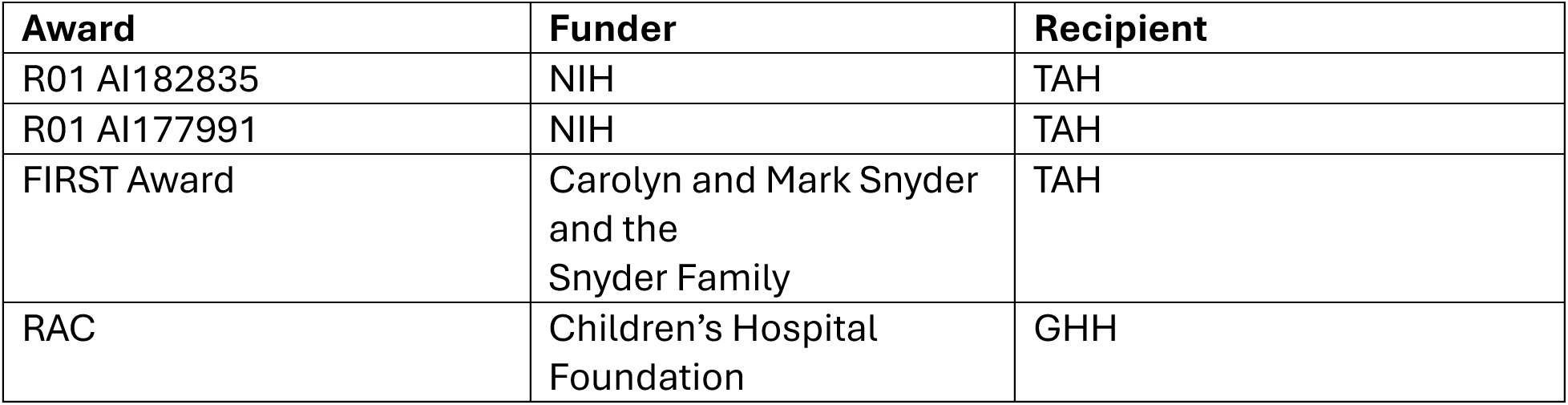

---

**Supplemental Data 1. Strains and DNA, including plasmids and oligonucleotides, used in this study**

**Supplemental Data 2. Transposon insertion hits recovered from positive selection screen under lethal heme conditions.** Table of the 18 transposon insertion sites identified among colonies surviving growth in 100 μM heme, listing gene name, locus tag, whether the gene is represented in the indexed transposon mutant library, and whether the hit met predefined inclusion criteria for follow-up validation as a candidate heme importer. The *cyd* locus (SAK_RS08800, highlighted) was selected for further study based on its predicted transmembrane topology and homology to the *E. coli* CydDC heme transporter.

**Supplemental Fig. 1:**
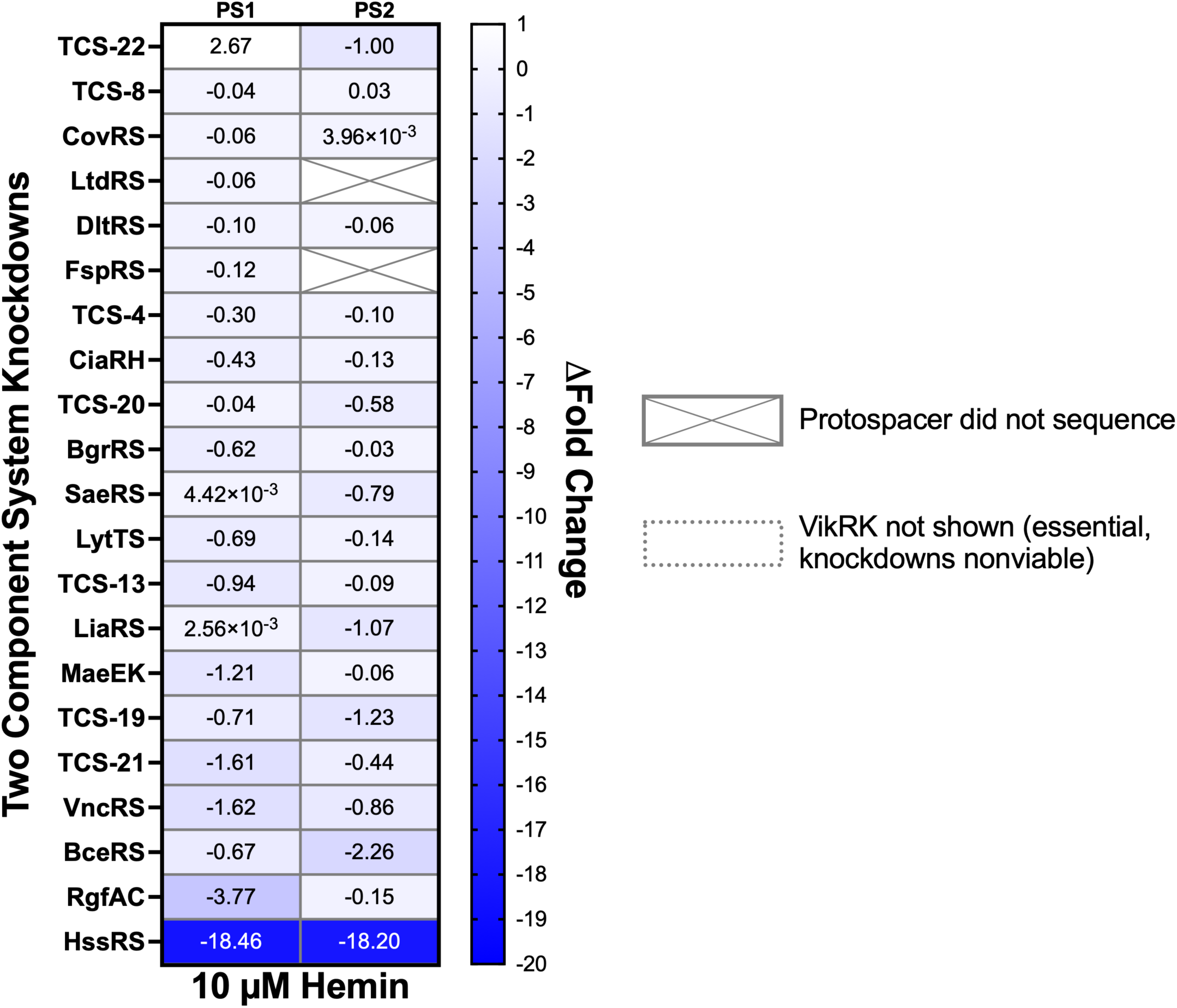
Fitness effects of hemin exposure on complete TCS CRISPRi library. Relative protospacer detection changes across nonselective TS broth and hemin exposure conditions for the CRISPRi library. Note that two intended knockdown strain protospacers (targeting LtdRS and FspRS systems) did not sequence and were not included in the analysis. Additionally, viable knockdowns of one essential TCS (VikRK) could not be isolated and so this system was not included in the analyses.

**Supplemental Fig. 2:**
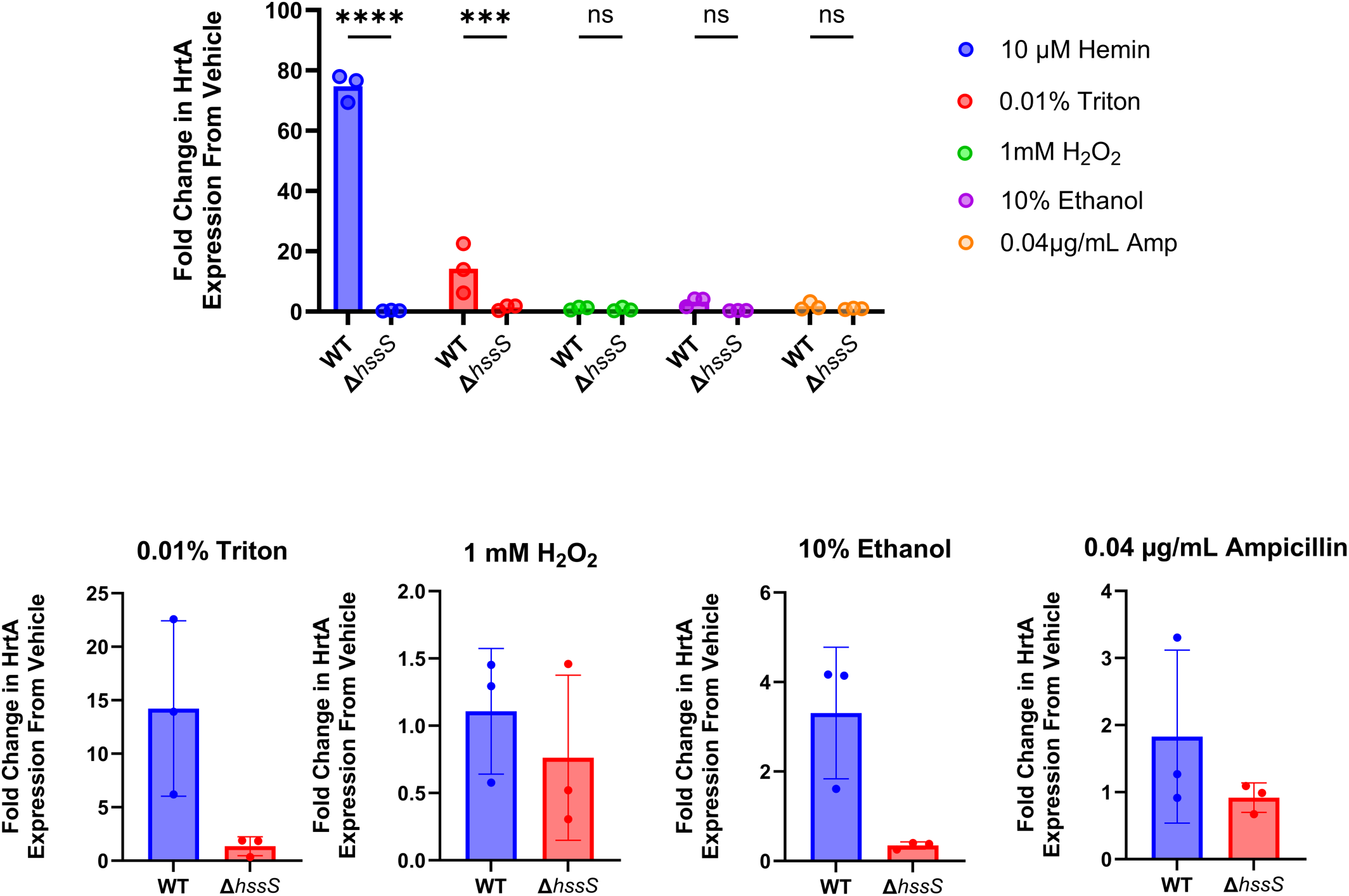
Heme uniquely induces HrtA expression through HssS signaling. RT-qPCR measurement of *hrtA* expression in WT and Δ*hssS* GBS following exposure to 10 μM hemin, 0.01% Triton X-100, 1 mM H₂O₂, 10% ethanol, or 0.04 μg/mL ampicillin. Fold change in *hrtA* expression is normalized to vehicle control. Bottom graphs are the same data as the top, zoomed in. n=3, **** p < 0.001, *** p < 0.005, ns = not significant, ANOVA with Bonferroni correction.

**Supplemental Fig. 3:**
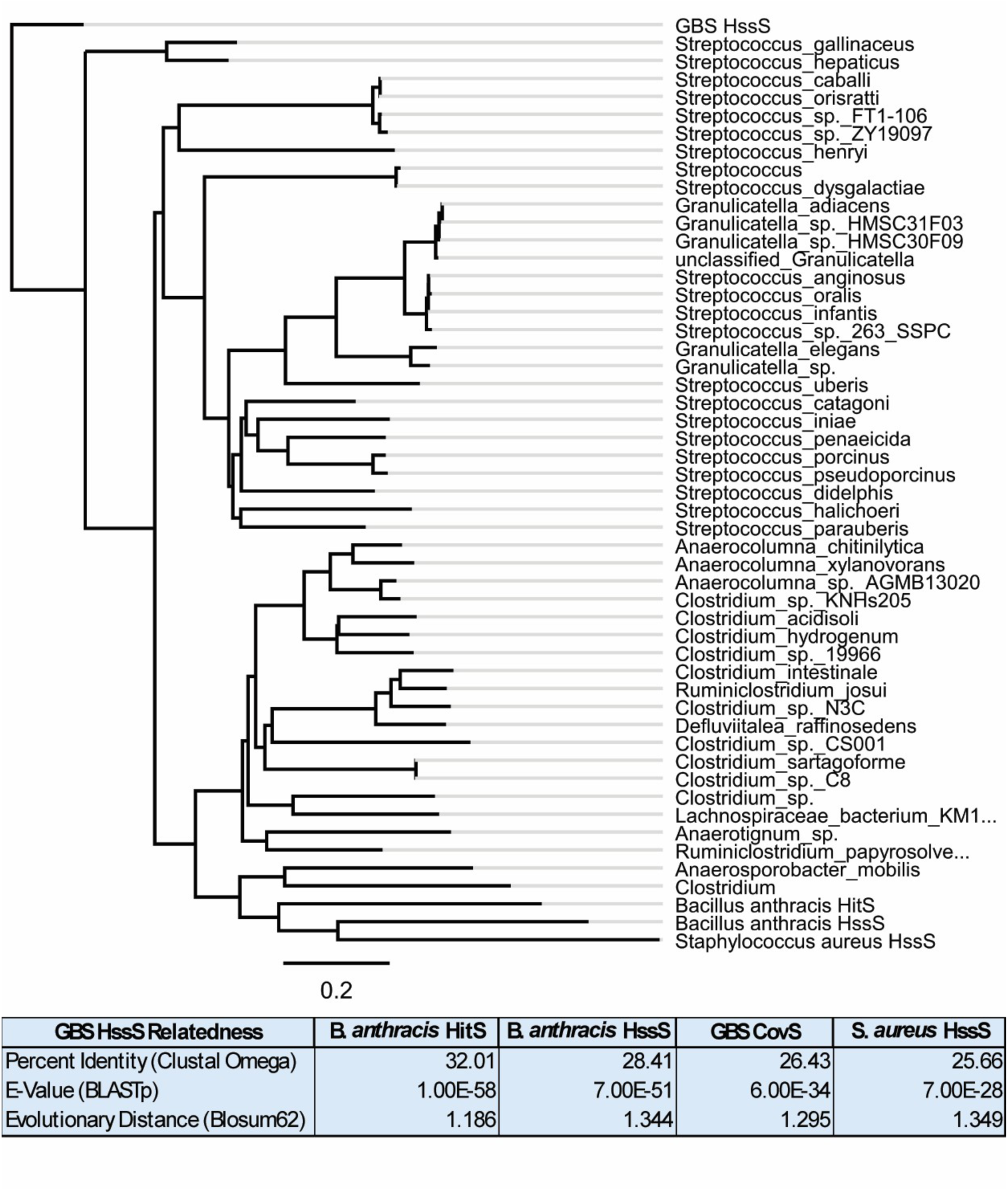
GBS HssS is phylogenetically divergent from characterized orthologs in *B. anthracis* and *S. aureus*. Phylogenetic tree of HssS protein sequences from GBS and closest HssS relatives from BLAST, with *B. anthracis* HitS, *B. anthracis* HssS, and *S. aureus* HssS included as outgroups. GBS HssS clusters within streptococcal relatives and is phylogenetically distant from the *B. anthracis* and *S. aureus* orthologs. Table below shows pairwise sequence relatedness metrics (percent identity by Clustal Omega, E-value by BLASTp, and evolutionary distance by Blosum62) between GBS HssS and each of the three outgroup sequences, as well as GBS CovS, a well-characterized GBS histidine kinase, for reference.

**Supplemental Fig. 4:**
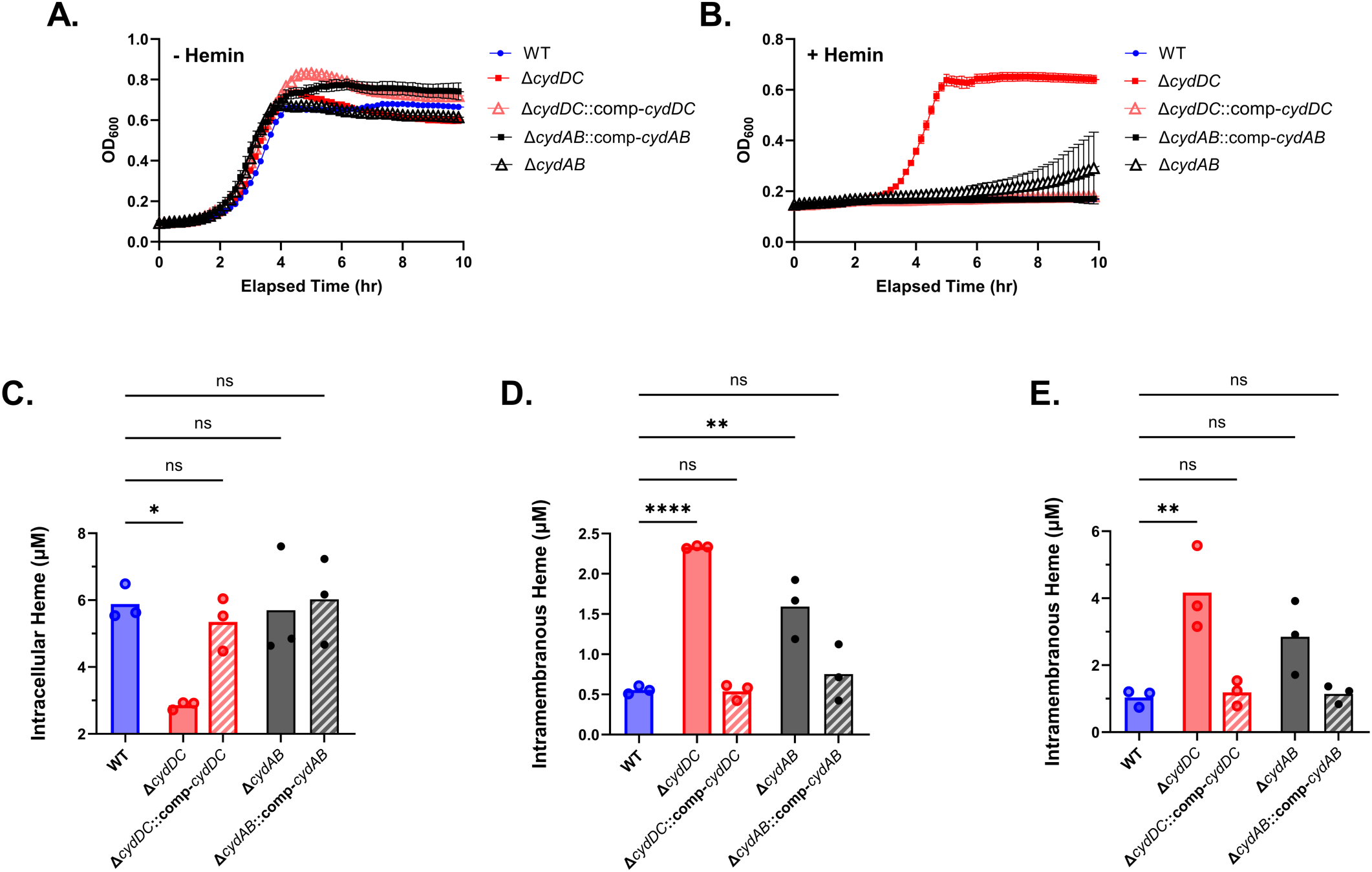
Loss of *cydDC*, but not *cydAB*, confers growth rescue under lethal hemin and does not alter intracellular heme concentrations. Growth curves of WT, Δ*cydDC*, Δ*cydDC*::comp-*cydDC*, Δ*cydAB*, and Δ*cydAB*::comp-*cydAB* in TS medium without hemin (**A.**) and with supplemental 10 µM hemin (**B**). Intracellular hemin concentrations after overnight night growth with supplemental 10 µM hemin (**C.**), intramembranous heme concentrations following growth in medium supplemented with 10 µM hemin (**D**) or 2.5% lysed whole blood (**E.**) are shown for all five strains. n=3 for all bar graphs, * p < 0.05, ** p < 0.01, **** p < 0.001, ns = not significant, ANOVA with Bonferroni correction.

**Supplemental Fig. 5:**
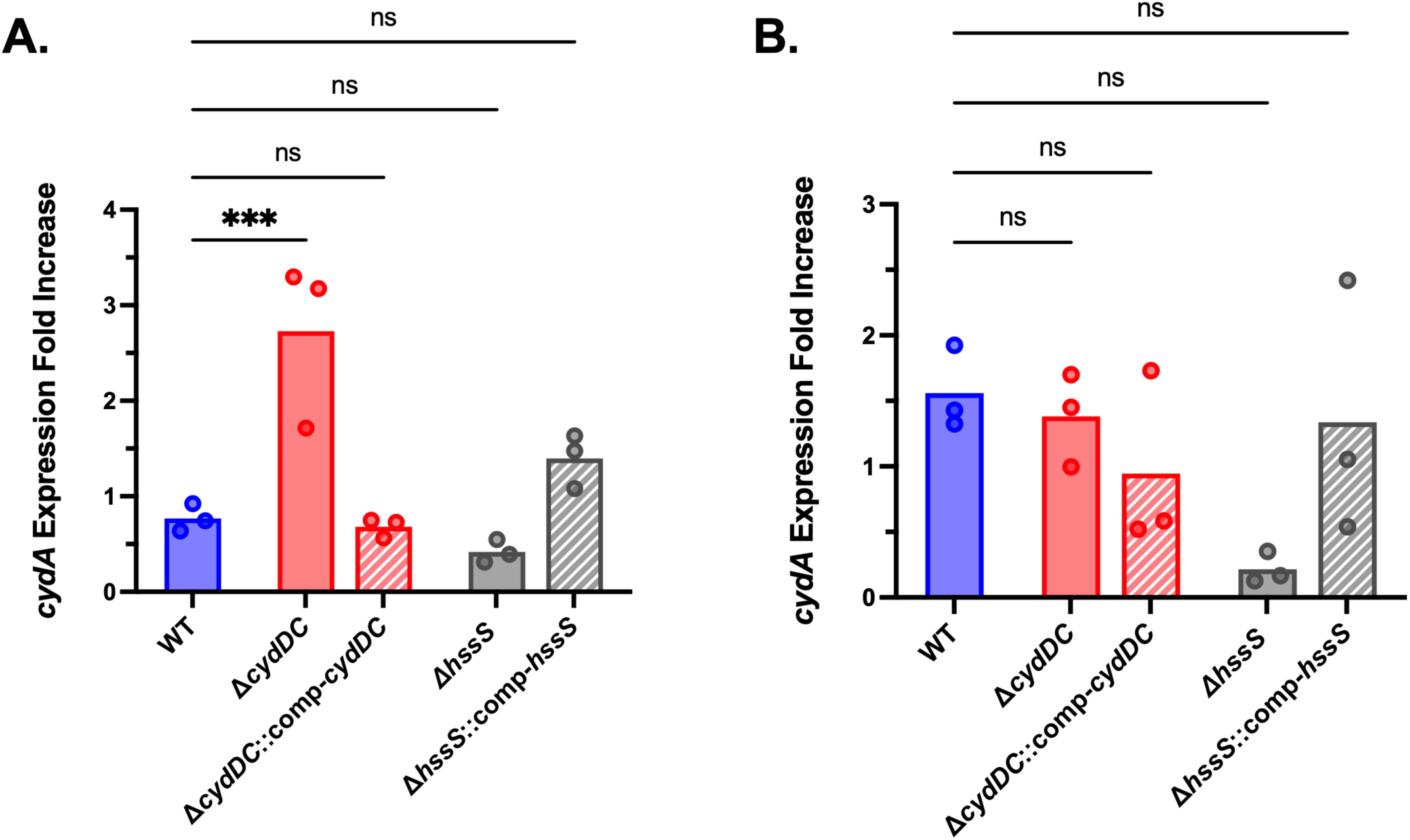
*cydA* expression during heme exposure. RT-qPCR measurement of *cydA* expression fold increase in WT, Δ*cydDC*, Δ*cydDC*::comp-*cydDC*, Δ*hssS*, and Δ*hssS*::comp-*hssS* strains following exposure to 10 µM hemin in TS medium (**A**) or 20% lysed human erythrocytes (**B**). Δ*cydDC* shows modestly increased *cydA* induction relative to WT under heme-supplemented conditions, These effects were substantially smaller in magnitude than HssRS-dependent regulation of the *hrtAB-hssRS* operon and are likely of little biological relevance. n=3, *** p < 0.005, ns = not significant, ANOVA with Bonferroni correction

**Supplemental Fig. 6:**
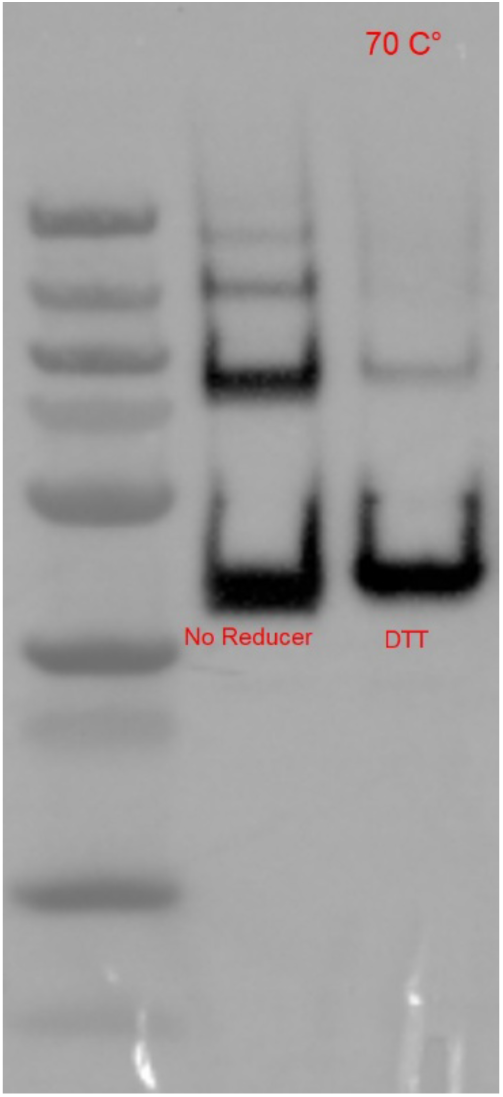
Recombinant HssS purification yields monomeric and dimeric protein under denaturing conditions. Western blot of polyhistidine-tagged recombinant HssS protein preparations resolved under denaturing, reducing SDS-PAGE conditions. Dominant bands consistent with HssS monomers are visible, along with a higher molecular weight band consistent with SDS-resistant HssS homodimers. The higher molecular weight bands diminished under harsher denaturing conditions (right band), consistent with its identity as a dimeric form of the protein.

**Supplemental Fig. 7:**
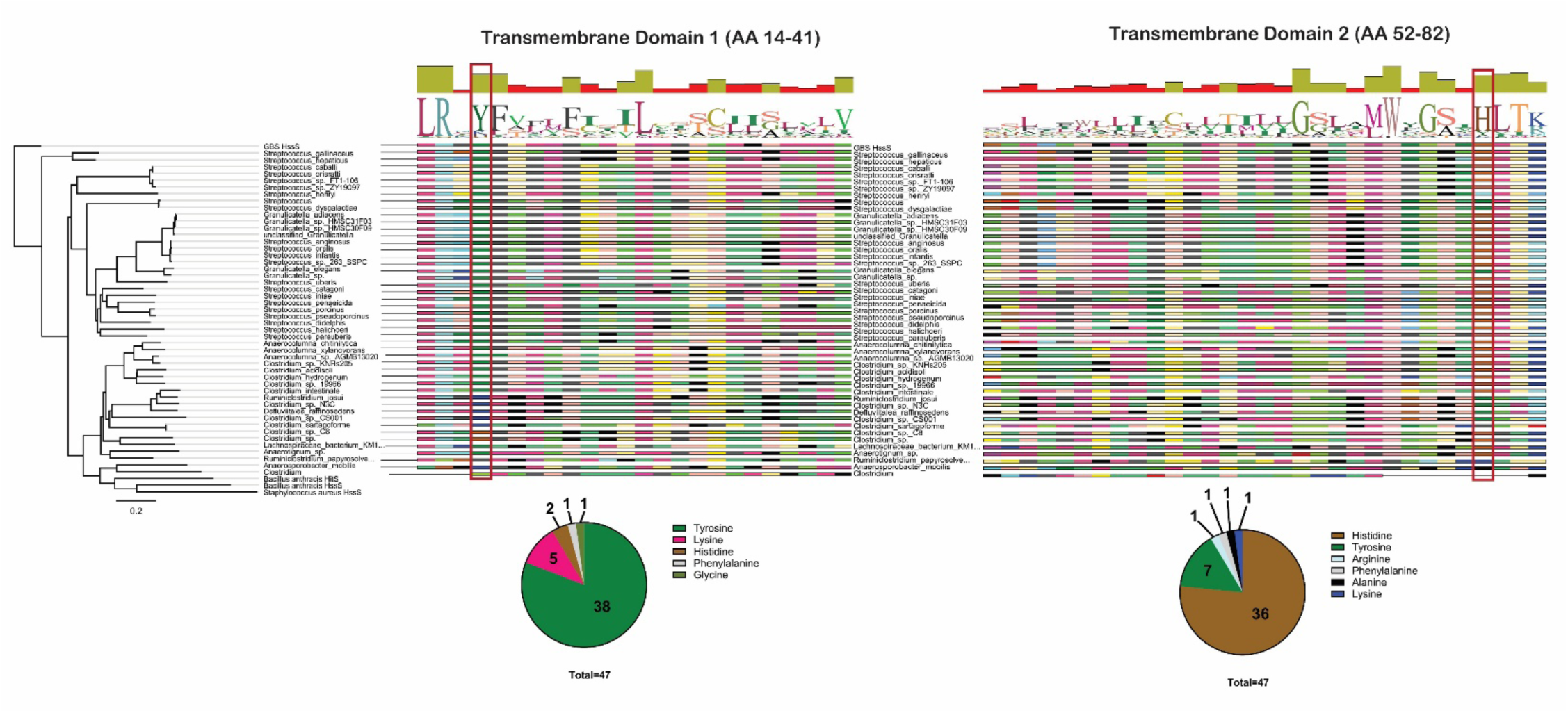
Conservation of putative heme-coordinating residues Y17 and H79 across HssS homologs. Multiple sequence alignment of HssS transmembrane domain 1 (AA 14–41, left) and transmembrane domain 2 (AA 52–82, right) from the 47 bacterial species in the Supplemental Figure 2 phylogenetic tree. Residue positions corresponding to GBS Y17 (TM1) and H79 (TM2) are highlighted with red boxes. Pie charts summarize the amino acid distribution at each highlighted position across all 47 sequences. Y17 is predominantly tyrosine (38/47), with minor representation of histidine, lysine, phenylalanine, and glycine. H79 is predominantly histidine (36/47), with moderate representation of tyrosine and minor representation of arginine, phenylalanine, alanine, and lysine, indicating conservation of both putative heme-coordinating residues across GBS HssS relatives.

**Supplemental Fig. 8:**
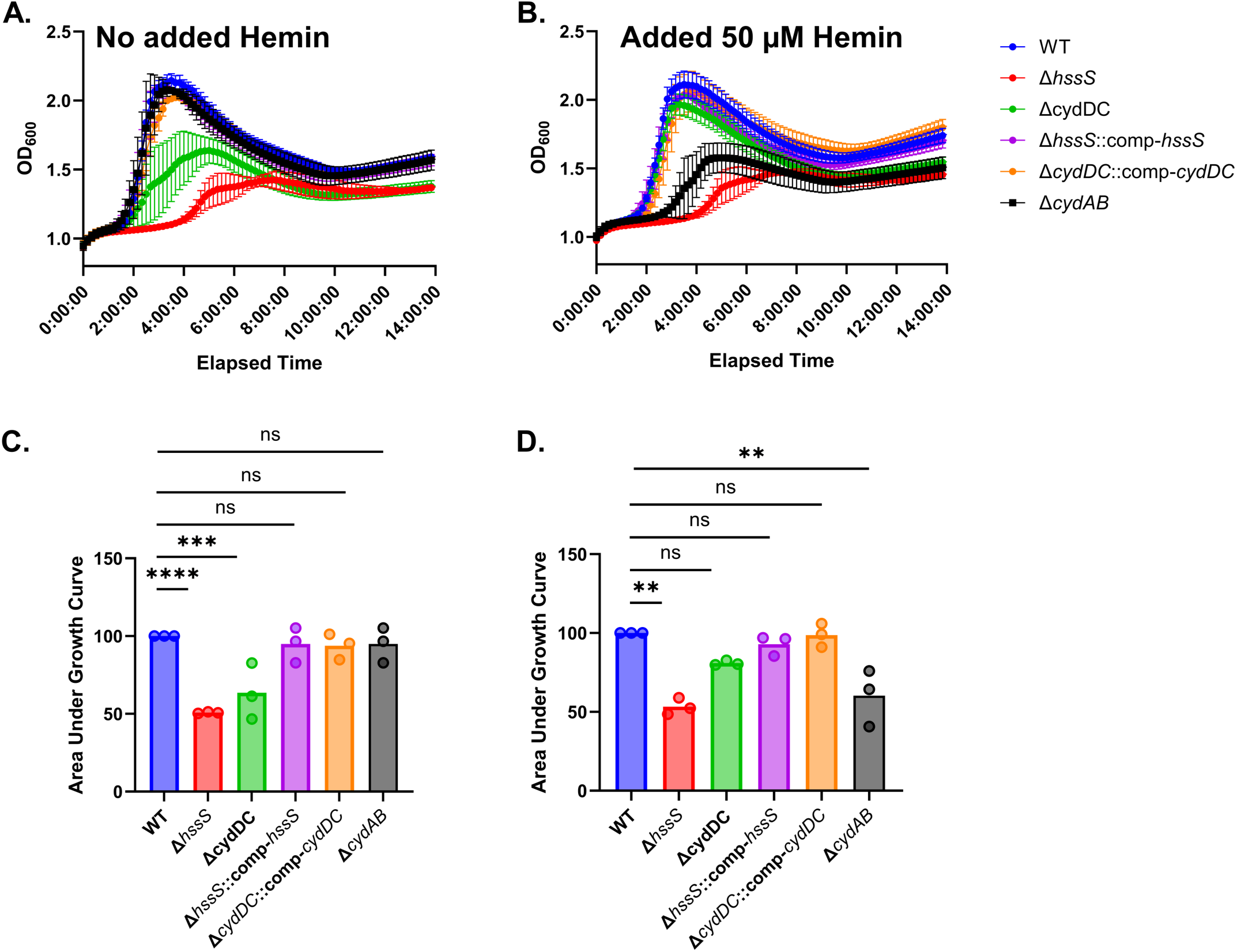
Loss of *cydAB* impairs growth in hemin-supplemented but not basal blood lysate, a phenotype opposite to that of Δ*cydDC*. Growth curves of WT, Δ*hssS*, Δ*cydDC*, Δ*hssS*::comp-*hssS*, Δ*cydDC*::comp-*cydDC*, and Δ*cydAB* in clarified human erythrocyte lysate without supplemental hemin (**A**) and with 50 μM supplemental hemin (**B**). Area under the growth curve comparisons for unsupplemented (**C**) and hemin-supplemented (**D**) conditions. Δ*cydAB* grows comparably to WT in unsupplemented lysate but is significantly impaired in hemin-supplemented conditions, a pattern opposite to that of Δ*cydDC*, which is impaired in unsupplemented lysate but recovers with excess heme. n=3, ** p < 0.01, *** p < 0.005, **** p < 0.001, ns = not significant, ANOVA with Bonferroni correction.

